# UBE3A–mediated p18/LAMTOR1 ubiquitination and degradation regulate mTORC1 activity and synaptic plasticity

**DOI:** 10.1101/242057

**Authors:** Jiandong Sun, Yan Liu, Yousheng Jia, Jennifer Tran, Xiaoning Hao, Weiju Lin, Gary Lynch, Michel Baudry, Xiaoning Bi

## Abstract

Accumulating evidence indicates that the lysosomal Ragulator complex is essential for full activation of the mechanistic target of rapamycin complex 1 (mTORC1). Abnormal mTORC1 activation has been implicated in several developmental neurological disorders, including Angelman syndrome (AS), which is caused by maternal deficiency of the ubiquitin E3 ligase UBE3A. Here we report that Ube3a regulates mTORC1 signaling by targeting p18, a subunit of the Ragulator. Ube3a ubiquinates p18, resulting in its proteasomal degradation, and Ube3a deficiency in hippocampus of AS mice results in increased lysosomal localization of p18 and other members of the Ragulator-Rag complex, such as RagA, and increased mTORC1 activity. P18 down-regulation by siRNA or shRNA in hippocampal CA1 neurons of AS mice reduces elevated mTORC1 activity and improves long-term potentiation (LTP) and dendritic spine maturation. Our results indicate that Ube3a-mediated regulation of p18 and subsequent mTORC1 signaling is critical for typical synaptic plasticity and dendritic spine development.

## Introduction

The mechanistic target of rapamycin (mTOR) is a highly conserved and ubiquitously expressed protein kinase complex, which plays important roles in cell survival, growth, and metabolism. mTOR, existing as mTORC1 and mTORC2, integrates extracellular signals (growth factors, neurotransmitters, nutrients, etc.) with intracellular energy levels and cellular stress status to regulate many important cellular functions (Laplante and Sabatini, 2012; Takei and Nawa, 2014). Not surprisingly, abnormal mTOR signaling has been implicated in various neurodevelopmental disorders and neuropsychiatric and neurological diseases (Costa-Mattioli and Monteggia, 2013). Recent evidence indicates that amino acid-induced lysosomal recruitment of mTORC1 is essential for its full activation (Jewell et al., 2013). In the presence of amino acids, mTORC1 is activated by binding to heterodimers consisting of Rag small guanosine triphosphatases (GTPases) RagA/B in their GTP-bound states and RagC/D in their GDP-bound states (Ham et al., 2016). Lysosomal localization of Rag dimers is maintained through their binding to the Ragulator complex, which consists of p18 (also known as LAMTOR1), p14 (LAMTOR2), MP1 (LAMTOR3), C7orf59 (LAMTOR4), and HBXIP (LAMTOR5) proteins (Bar-Peled et al., 2012; Nada et al., 2009; Sancak et al., 2010); acylation of p18 is essential for anchoring the Ragulator complex to endosomal/lysosomal membranes (Nada et al., 2009). Although it has been shown that lysosomal localization and interaction with Rag GTPases are essential for p18 to regulate mTORC1 activation (Sancak et al., 2010), little is known regarding the regulation of p18 levels. A previous study combining single-step immuno-enrichment of ubiquitinated peptides with high-resolution mass spectrometry revealed that p18 was ubiquitinated at residues K20 and K31 in HEK cells and MV4-11 cells (Wagner et al., 2011). However, the E3 ligase responsible for p18 ubiquitination was not identified.

UBE3A, an E3 ligase in the ubiquitin-proteasomal system, plays important roles in brain development and normal function, as UBE3A deficiency results in Angelman syndrome (AS) (Williams et al., 1990), while UBE3A over-expression increases the risk for autism (Cook et al., 1997). We recently reported that imbalanced signaling of the mTOR pathway, with increased mTORC1 and decreased mTORC2 activation, plays important roles in the motor dysfunction and abnormal dendritic spine morphology of Purkinje neurons in AS mice (Sun et al., 2015a). A similar abnormal mTOR signaling is critically involved in Ube3a deficiency-induced impairment in hippocampal synaptic plasticity and fear-conditioning memory (Sun et al., 2016). Furthermore, inhibition of mTORC1 by rapamycin treatment not only reduced mTORC1 activity but also normalized mTORC2 activity, suggesting that mTORC1 overactivation is the trigger for alterations in mTOR signaling in AS mice. However, how Ube3a deficiency results in mTORC1 over-activation remains unknown. Considering the critical roles of p18 in mTORC1 activation, we investigated the potential regulation of p18 levels by Ube3a. Here, we demonstrate that Ube3a directly ubiquitinates p18 and targets it for proteasomal degradation, which contributes to the regulation of mTORC1 signaling and activity-dependent synaptic remodeling. In the absence of Ube3a, p18 accumulates in neurons, resulting in mTORC1 overactivation, abnormal synaptic morphology, and impaired synaptic plasticity. These findings reveal a previously unidentified regulatory mechanism for mTORC1 activation and suggest potential therapeutic targets for cognitive disorders associated with abnormal mTORC1 signaling.

## Results

### 1. P18 is a Ube3a substrate

Although it has been shown that p18 plays essential roles in mTOR and MAP kinase signaling and other cell functions, very little is known regarding its biosynthesis and degradation. We first determined whether Ube3a could regulate p18 levels in heterologous cells. Western blot analysis showed that Ube3a knockdown in COS-1 cells by siRNA resulted in increased p18 levels, as compared to scrambled control siRNA (Figure 1A). Sequence analysis revealed the presence of 6 lysine residues in p18, which could represent ubiquitination sites (Figure 1B). To assess whether p18 could be a Ube3a substrate, we first determined whether these 2 proteins exhibited direct interactions. Co-immunoprecipitation experiments using extracts from COS-1 cells transfected with Ube3a and Flag-p18 showed that p18 could bind to Ube3a in an E3 ligase activity-independent manner, since p18 could also bind to an inactive form of Ube3a with a mutation in the catalytic site, Ube3a-C833A (Kumar et al., 1999) (referred to as ∆Ube3a hereafter) (Figure 1C). *In vitro* ubiquitination assays using purified recombinant p18 and a Ube3a ubiquitination assay kit showed that p18 ubiquitination was only observed in the presence of Ube3a and ubiquitin, E1, E2, and ATP (Figure 1D). We then determined whether Ube3a could ubiquitinate p18 in intact cells using His-ubiquitin pull-down assay. COS-1 cells were co-transfected with p18 and His-ubiquitin plus either an empty vector, or Ube3a or ∆Ube3a. Ubiquitinated proteins were extracted by Co^2+^-affinity chromatography and analyzed by Western blot. Co-transfection with Ube3a, but not ∆Ube3a, resulted in massive p18 ubiquitination (Figure 1E and Figure 1-figure supplement 1A). In addition, transfection with Ube3a, but not ∆Ube3a, resulted in decreased p18 levels (Figure 1F and Figure 1-figure supplement 1B), indicating that Ube3a-mediated regulation of p18 levels depends on its E3 ligase activity and p18 ubiquitination. Finally, to confirm that Ube3a-mediated p18 ubiquitination was on lysine residues, we simultaneously mutated all lysine residues into arginine (∆K) in Flag-p18. COS-1 cells were first transfected with either Ube3a siRNA or a scrambled control siRNA, followed by transfection with Flag-p18 or Flag-p18∆K and His-ubiquitin. His-ubiquitin pull-down assay showed that levels of p18-immunopositive high molecular weight bands, i.e., ubiquitinated, were significantly reduced following Ube3a siRNA knockdown (Figure 1G and Figure 1-figure supplement 1C), and the degree of reduction (100.0 ± 1.8 vs 66.8 ± 3.7, n = 4, p < 0.001) corresponded to the extent of Ube3a down-regulation (100.0 ± 2.4 vs 60.9 ± 5.2, n = 4, p < 0.001). Furthermore, p18 ubiquitination was abolished by K-R mutations (Figure 1G and Figure 1-figure supplement 1C). ∆K mutations did not significantly affect the interaction between p18 and Ube3a (Figure 1C). These results confirmed that p18 is ubiquitinated by Ube3a at lysine residues.

**Figure 1.**
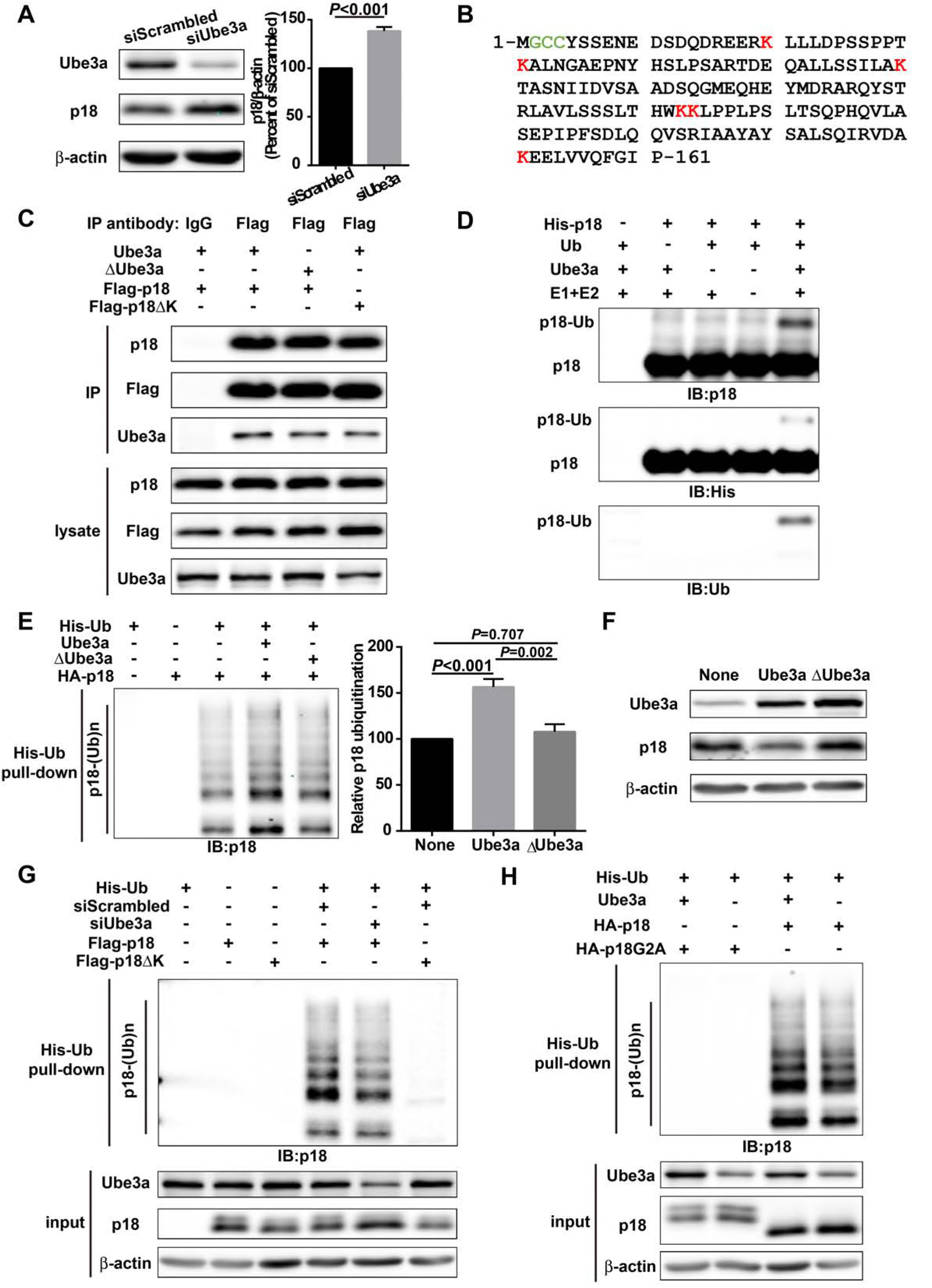
P18 is a Ube3a substrate. (A) Western blot analysis using anti-Ube3a, p18, or β-actin antibodies of lysates from COS-1 cells transfected with scrambled siRNA or Ube3a siRNA. Right, quantitative analysis of blots. N = 6, p < 0.001 (unpaired, two-tailed Student’s t test). (B) Amino acid sequence of human p18. G2 is a myristoylation site. C3 and C4 are palmitoylation sites. K20, K31, K60, K103, K104, and K151 are potential ubiquitination sites. (C) Interaction between p18 and Ube3a. Lysates from COS-1 cells transfected with the indicated cDNAs in expression vectors were immunoprecipitated with an anti-Flag antibody or control IgG and probed with the indicated antibodies. The presence of Flag-p18 in precipitates was confirmed with anti-p18 and anti-Flag antibodies. (D) *In vitro* ubiquitination of p18 by recombinant Ube3a. Reaction products were analyzed by Western blots with p18, His, and ubiquitin antibodies. Note that the p18-Ub band is present only when all reaction elements were added. (E) Overexpression of Ube3a, but not ∆Ube3a, enhances p18 ubiquitination in COS-1 cells. His-tagged ubiquitinated proteins in cells co-transfected with HA-p18 plus empty vectors, wild-type Ube3a (Ube3a), or its inactive form Ube3a-C833A (∆Ube3a) were precipitated using Talon resin and probed with anti-p18 antibodies. Ubiquitinated p18 proteins are labeled with “p18-(Ub)n”. Right, quantification of the relative abundance of ubiquitinated p18 (means ± SEM, p < 0.001 None vs Ube3a, p = 0.002 Ube3a vs ∆Ube3a, p = 0.707 None vs ∆Ube3a, n = 3, one-way ANOVA with Tukey’s post hoc analysis). (F) Western blot analysis using anti-Ube3a, p18 or β-actin antibodies on lysates from COS-1 cells transfected with empty vector (None), Ube3a, or ∆Ube3a vectors. (G) siRNA knock-down of Ube3a in COS-1 cells reduces p18 ubiquitination. COS-1 cells were incubated with Ube3a siRNA or scrambled control siRNA 48 h before transfection with Flag-p18 or Flag-p18∆K and His-ubiquitin. Twenty-four h later, ubiquitinated proteins were isolated by Co^2+^-affinity chromatography. Levels of ubiquitinated p18 protein (p18-(Ub)n, upper panel) were determined by Western blots. Levels of input proteins were also evaluated by Western blots probed with Ube3a, p18, and β-actin antibodies (lower panel). (H) His-ubiquitin pull down assay performed using HA-p18 or HA-p18G2A. Upon purification, levels of ubiquitinated p18 (upper panel) were determined by Western blot analysis. Lower panel, input of Ube3a, p18, and β-actin. See also Figure 1-figure supplement 1.

Previous studies have revealed that p18 is anchored to lysosomal membranes through myristate and palmitate modifications at G2 and C3/C4, respectively (Nada et al., 2014). We confirmed that wild-type (WT) p18 was indeed localized at the lysosomal surface, while its myristoylation-defective mutant, p18G2A, failed to localize to the lysosomal surface (Figure 1-figure supplement 1D), and nearly completely lost its ability to be ubiquitinated (Figure 1H and Figure 1-figure supplement 1E), suggesting that myristoylation-dependent lysosomal localization of p18 is required for Ube3a-mediated p18 ubiquitination.

### 2. P18 is essential for lysosomal localization of Ragulator and RagGTPases in hippocampal neurons

Since there is little information regarding p18 in the central nervous system (CNS), we next characterized p18 expression in hippocampal neurons. Double immunolabeling with antibodies against p18 and LAMP2, a well-characterized lysosomal marker (Eskelinen, 2006), showed that p18 was co-localized with LAMP2, not only in cell bodies but also in dendrites of cultured mouse hippocampal neurons (Figure 2A). To determine whether p18 could interact with other members of the Ragulator complex in neurons as in other cell types, lysates from cultured hippocampal neurons were immunoprecipitated with a p18 antibody and the precipitated material was probed with anti-p18, anti-p14, or anti-MP1 antibodies. P18, p14, and MP1 were detected in anti-p18-but not in control IgG-pull-down proteins (Figure 2B). Immunoprecipitates prepared with a RagA but not a control IgG antibody from hippocampal lysates of WT mice consistently contained RagA, RagB, RagC, p18, and p14 (Figure 2B). These results indicate that p18 interacts with other members of the Ragulator complex, and that the Ragulator interacts with Rag GTPases in hippocampal neurons, as in other cell types.

**Figure 2.**
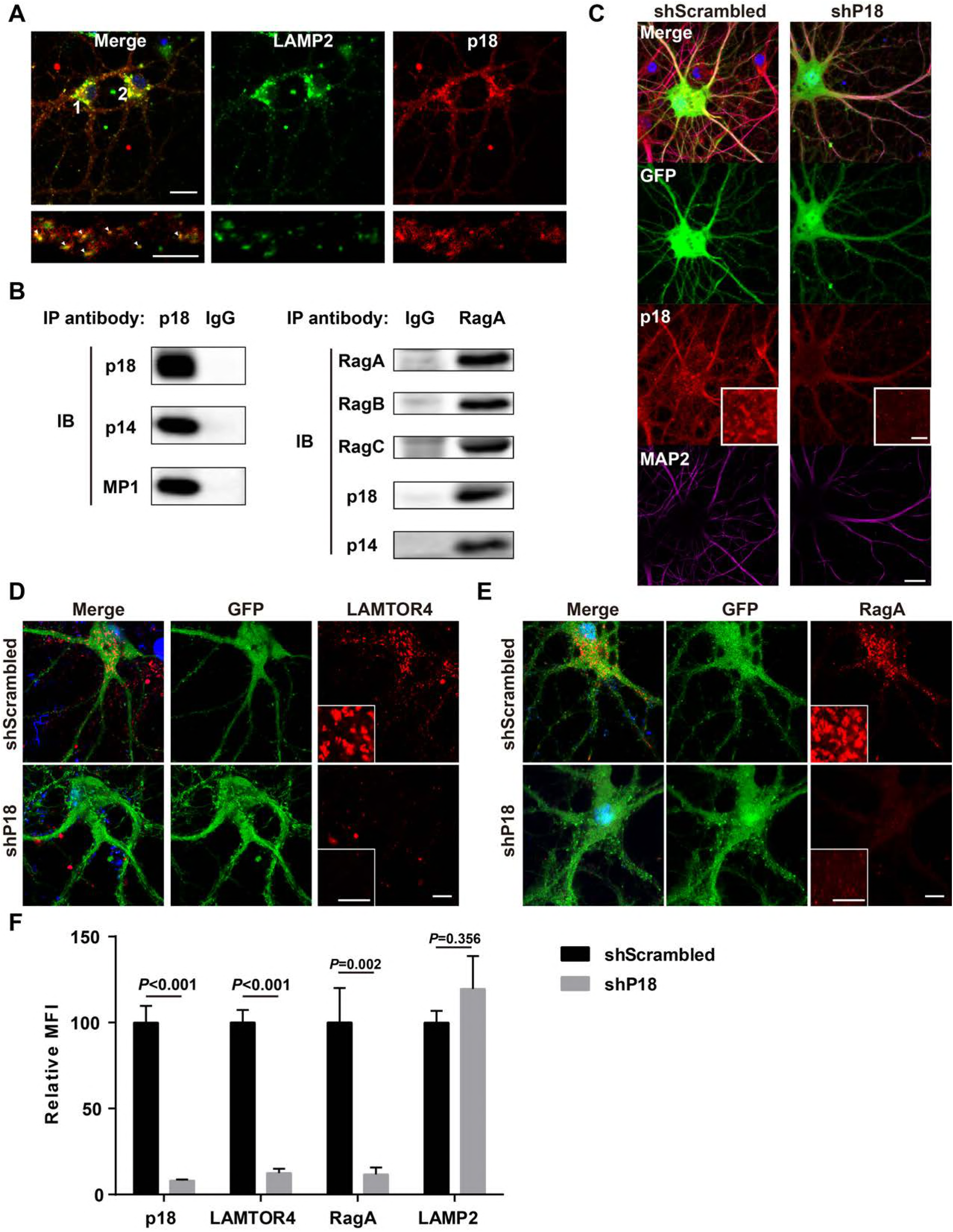
Characterization of p18 in hippocampal neurons. (A) Images of cultured hippocampal neurons co-immunostained for lysosomal protein LAMP2 (green) and p18 (red). Insets are enlarged images of LAMP2- and p18-immunoreactive puncta along the dendrites. Arrowheads indicate co-localized puncta. Scale bar: top, 20 μm; inset, 10 μm. (B) Left, p18 forms complex with p14 and MP1 in hippocampal neurons. Lysates from cultured hippocampal neurons were immunoprecipitated with an anti-p18 antibody or control IgG and probed with the indicated antibodies. Right, RagA co-immunoprecipitates RagB, RagC, p18, and p14. Lysates from mouse hippocampi were immunoprecipitated with an anti-RagA antibody or control IgG and probed with the indicated antibodies. (C) Images of cultured hippocampal neurons co-immunostained for p18 (red) and MAP2 (magenta). Neurons were infected with shRNA AAV directed against p18 with GFP co-expression or scrambled shRNA control before processing for immunofluorescence assay and imaging. Scale bar, 20 μm; inset, 5 μm. (D) Images of hippocampal neurons stained for LAMTOR4 (red). Cells were infected and processed as in (C). Scale bar, 10 μm; inset, 5 μm. (E) Images of hippocampal neurons stained for RagA (red). Cells were infected and processed as in (C). Scale bar, 10 μm; inset, 5 μm. (F) Quantification of fluorescent signals for p18 (n = 13, p < 0.001), LAMTOR4 (n = 11, p < 0.001), RagA (n = 6, p = 0.002), and LAMP2 (n = 6, p = 0.356) in control shRNA and p18 shRNA-transfected neurons. Student’s t test. See also Figure 2-figure supplement 1.

To further investigate whether p18 also serves as an anchor for the Ragulator-Rag complex in the brain, neurons were infected with shRNA AAV directed against p18 with GFP co-expression in order to decrease p18 expression, and the resulting effects on the Ragulator-Rag complex were determined. Confocal images of transfected neurons indicated that p18 shRNA transfection efficiently reduced p18 expression in cultured neurons (Figure 2C,F). Notably, levels of lysosome-localized LAMTOR4 and RagA were significantly reduced following p18 shRNA knockdown in neurons (Figure 2D-F and Figure 2-figure supplement 1A,B), while lysosomal morphology was not obviously affected (Figure 2F and Figure 2-figure supplement 1C). Thus, p18 is required for lysosomal targeting of the Ragulator-Rag complex in neurons.

### 3. Ube3a regulates p18 levels in a proteasome-dependent manner in hippocampal neurons

We then determined whether Ube3a deficiency in neurons could result in increased p18 levels using AS mice. Western blot results showed that p18 levels were markedly increased in crude membrane fractions (P2) of hippocampus from AS mice, as compared to WT mice (Figure 3A and Figure 3-figure supplement 1A), while there was no significant change in levels of p14, MP1, as well as Rag GTPases (Figure 3A and Figure 3-figure supplement 1A). Importantly, although rapamycin treatment normalized mTORC1 and mTORC2 signaling (Sun et al., 2016), it did not reverse the increase in p18 levels in AS mice (Figure 3-figure supplement 1A, same samples as in Sun et al., 2016). These results suggest that increased p18 levels in AS mice are independent of mTORC1 activity, and that increased mTORC1 activity might be downstream of p18 level increases.

**Figure 3.**
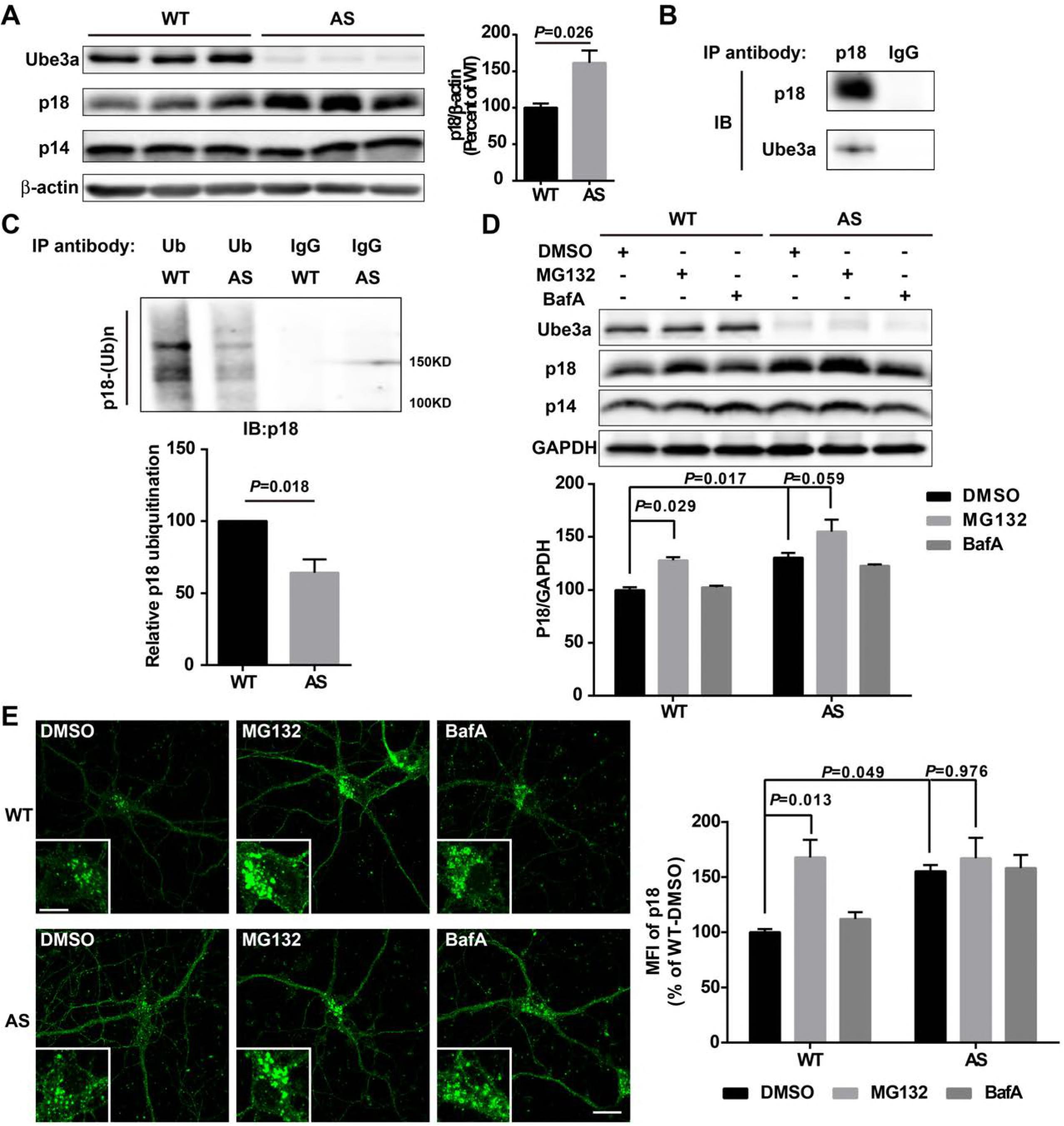
Ube3a regulates p18 levels in a proteasome-dependent manner in hippocampal neurons. (A) Left, Western blot analysis of p18 and p14 levels in crude membrane fractions (P2) of hippocampi from WT and AS mice. Right, quantitative analysis of blots. Results are expressed as % of values in WT mice and shown as means ± S.E.M. N = 3, p = 0.026 (unpaired, two-tailed Student’s t test). (B) Interactions between Ube3a and p18 in hippocampal neuron cultures. Western blot analysis with anti-p18 and -Ube3a antibodies of immunoprecipitation performed with anti-p18 antibodies or control IgG. (C) Immunoprecipitation of hippocampal P2 fractions from WT and AS mice under denaturing conditions was performed with anti-ubiquitin antibodies or control IgG and Western blots were labelled with anti-p18 antibodies. Ubiquitinated p18 proteins are indicated as “p18-(Ub)n”. Lower panel: quantification of the relative abundance of ubiquitinated p18 in hippocampus of WT and AS mice (mean ± SEM, p = 0.018 compared to WT mice, n = 3, Student’s t test). (D) Effects of acute MG132 or Bafilomycin A1 (BafA) treatment on p18 and p14 levels in hippocampus slices of WT and AS mice. Upper panel: representative Western blot images; lower panel: quantitative analysis of blots in upper panel. N = 3, p = 0.029 WT/DMSO vs WT/MG132, p = 0.017 WT/DMSO vs AS/DMSO, p = 0.059 AS/DMSO vs AS/MG132, two-way ANOVA with Tukey’s post-test. (E) Representative images of p18 in WT and AS hippocampal neurons treated with DMSO, MG132, and BafA; insets: enlarged cell bodies. Right: Quantitative analysis of images. Data are expressed as mean ± SEM. N = 3, p = 0.013 WT/DMSO vs WT/MG132, p = 0.049 WT/DMSO vs AS/DMSO, p = 0.976 AS/DMSO vs AS/MG132; two-way ANOVA with Tukey’s post hoc analysis. Scale bar = 20 μm and 10 μm in insets. See also Figure 3-figure supplement 1.

Our data suggested that increased p18 levels in hippocampus of AS mice could be due to the lack of Ube3a-mediated p18 ubiquitination and subsequent degradation. To further confirm this possibility, we first determined whether p18 could bind to Ube3a in hippocampal neuronal cultures (Figure 3B). Ubiquitinated proteins from hippocampal neuronal cultures or P2 fractions of hippocampi from WT and AS mice were immunoprecipitated with ubiquitin antibodies under denaturing conditions, and precipitated proteins were processed for Western blot with ubiquitin and p18 antibodies. Both p18 and ubiquitin antibodies labeled high molecular weight bands, and the intensity of p18-immunopositive bands was much weaker in samples from AS mice than WT mice (Figure 3C and Figure 3-figure supplement 1B,C), indicating that the increase in p18 levels in AS mice was likely due to a deficit in Ube3a-mediated p18 ubiquitination and degradation.

To determine whether Ube3a-mediated regulation of p18 was proteasome- and/or lysosome-dependent, acute hippocampal slices from WT and AS mice were treated with either a proteasome inhibitor, MG132 (10 μM), or a lysosome inhibitor, the vacuolar H^+^-ATPase (V-ATPase) inhibitor, bafilomycin A1 (BafA, 100 nM), for 30 min. These concentration and treatment duration have previously been shown to significantly inhibit proteasome or lysosomal function, respectively (Kim et al., 2015). As expected, levels of p18 were significantly higher in vehicle-treated AS slices than in vehicle-treated WT slices. Incubation of hippocampal slices with MG132, but not BafA, significantly increased p18 levels in WT slices and marginally in AS slices (Figure 3D). In addition, MG132 treatment (10 μM, 4 h) markedly increased p18 intensity in WT neuronal cultures, while BafA treatment (100 nM, 4 h) only slightly increased p18 in both WT and AS neuronal cultures (Figure 3E). These results strongly suggest that Ube3a decreases p18 levels via ubiquitination followed by proteasomal degradation.

### 4. Increased p18 levels in AS mice are associated with increased lysosomal localization of Ragulator-Rag complex and mTOR

Immunofluorescent staining showed that p18 was clearly co-localized with LAMP2 in CA1 pyramidal neurons, especially in cell bodies, in both WT and AS mice, and more p18/LAMP2 double-stained puncta were detected in AS mice than in WT mice (Figure 4A,B). Similarly, lysosomal localization of other members of the Ragulator, LAMTOR4 (Figure 4B and Figure 4-figure supplement 1A), p14 and MP1 (Figure 4-figure supplement 1D) was also increased in CA1 pyramidal cell bodies of AS, as compared to WT mice. Furthermore, dual immunohistochemical staining for either RagA/B or mTOR with LAMP2 showed that co-localization of these proteins with LAMP2 was markedly increased in AS mice, as compared to WT mice (Figure 4B and Figure 4-figure supplement 1B-D). P18 was also clearly co-localized with LAMP2 in apical dendrites in hippocampal CA1 region of adult mice, and more p18/LAMP2 double-stained puncta were detected in AS than in WT mice (Figure 4D and Figure 4-figure supplement 1E). Similarly, more mTOR proteins were co-localized with LAMP2 in CA1 apical dendrites of AS than WT mice (Figure 4C,D). We next evaluated the co-localization of p-mTOR, the active isoform of mTOR, with LAMP2. More p-mTOR/LAMP2 double-stained puncta were observed in both soma and apical dendrites in hippocampal CA1 region of AS than in WT mice (Figure 4E,F and Figure 4-figure supplement 1F).

**Figure 4.**
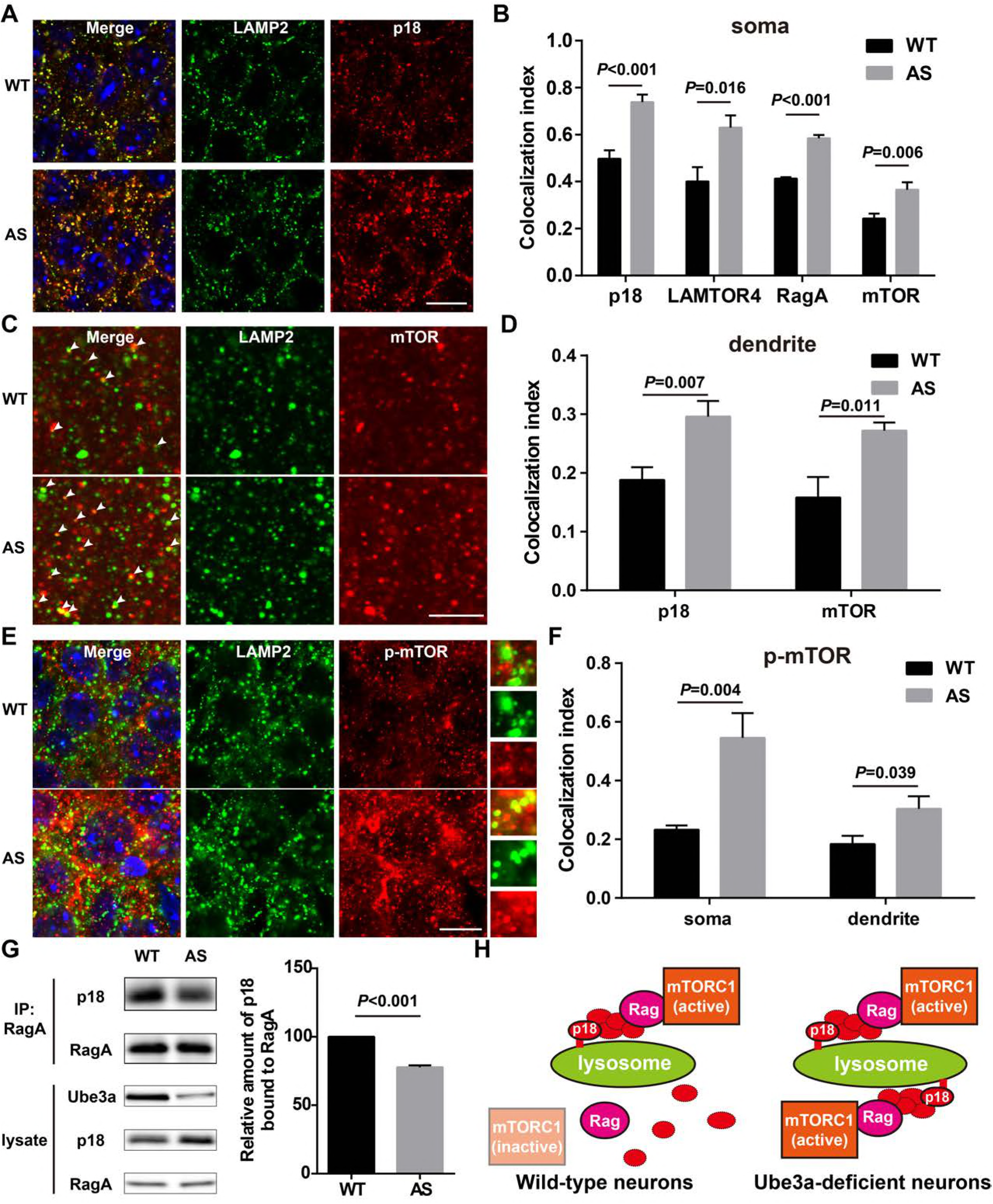
Lysosomal localization of Ragulator-Rag complex and mTOR/p-mTOR in WT and AS mice. (A) Co-localization of p18 (red) with LAMP2 (green) in cell bodies of CA1 pyramidal neurons from WT and AS mice. Scale bar = 10 μm. (B) Quantification of p18-LAMP2 (n = 8, p < 0.001), LAMTOR4-LAMP2 (n = 6, p = 0.016), RagA-LAMP2 (n = 6, p < 0.001), and mTOR-LAMP2 (n = 8, p = 0.006) colocalization in cell bodies of CA1 pyramidal neurons from WT and AS mice. Unpaired t-test. (C) Representative images of apical dendrites of CA1 pyramidal neurons stained with anti-mTOR (red) and -LAMP2 (green) antibodies. Arrowheads indicate puncta with dual staining. Scale bar = 5 μm. (D) Quantification of p18-LAMP2 (n = 8, p = 0.007) and mTOR-LAMP2 (n = 7, p = 0.011) co-localization in apical dendrites of CA1 pyramidal neurons from WT and AS mice. Unpaired t-test. (E) Co-localization of p-mTOR (red) with LAMP2 (green) in cell bodies of CA1 pyramidal neurons from WT and AS mice. Scale bar = 10 μm. Insets show selected fields that were magnified ten times. (F) Quantification of p-mTOR-LAMP2 co-localization in cell bodies (p = 0.004) and dendrites (p = 0.039) of CA1 pyramidal neurons from WT and AS mice. N = 6, unpaired t-test. (G) Homogenates from WT and AS mouse hippocampus were immunoprecipitated with an anti-RagA antibody and probed with the indicated antibodies. Right, quantification of the relative abundance of p18 bound to RagA (mean ± SEM, p < 0.001, n = 3, Student’s t test). (H) Model proposing that the Ragulator interacts with Rag, which in turn recruits mTORC1 to be activated on lysosomes in neurons. In Ube3a-deficient neurons, increased Ragulator-Rag complex on lysosomes results in mTORC1 over-activation. See also Figure 4-figure supplement 1.

In addition to providing a platform for recruiting Rag GTPases and subsequently mTORC1 to lysosomes, p18 has also been shown to function as a RagA/B GEF, which facilitates the exchange of GDP from RagA/B to GTP (Bar-Peled et al., 2012). To test whether increased p18 levels in AS mice could lead to increased levels of GTP-bound RagA/B, we used the widely-used co-immunoprecipitation assay of p18 and RagA, based on the observation that GTP-bound RagA/B has a lower affinity for the Ragulator, as compared to GDP-bound RagA/B (Bar-Peled et al., 2012; Castellano et al., 2017). Co-immunoprecipitation results showed that levels of p18 immunoprecipitated by RagA antibodies were significantly lower in samples from AS mice, as compared to WT mice (Figure 4G). Collectively, these results showed that increased p18 levels in hippocampus of AS mice facilitate lysosomal anchoring of the Ragulator-Rag complex and the activation of RagA/B and mTORC1 (see schematic in Figure 4H).

### 5. P18 downregulation counteracts Ube3a deficiency-induced abnormal mTOR signaling and changes in dendritic spine morphology and actin polymerization in cultured hippocampal neurons

We next directly tested whether reducing p18 levels could ameliorate Ube3a deficiency-induced mTORC1 overactivation in cultured hippocampal neurons. P18 expression was reduced by neuronal transfection with a set of p18 shRNA lentiviruses, while Ube3a knockdown was achieved with Accell Ube3a siRNA (Figure 5A,B). Ube3a down-regulation resulted in increased p18 levels (Figure 5A,B), in parallel with increased mTORC1 activation, as reflected by phosphorylation of mTOR and its downstream substrate S6, and decreased mTORC2 activation, as reflected by decreased (p-)PKCα levels. Increased mTORC1 activation and decreased mTORC2 activation were reversed by p18 knockdown (Figure 5A,B). These results strongly suggest that increased p18 levels contribute to Ube3a deficiency-induced abnormal mTOR signaling. Importantly, p18 shRNA knockdown in the absence of Ube3a knockdown led to a reduction in mTORC1 activation and a concomitant increase in the activity of mTORC2 (Figure 5A,B), which further underscores the notion that mTOR signaling is very sensitive to changes in p18 levels.

**Figure 5.**
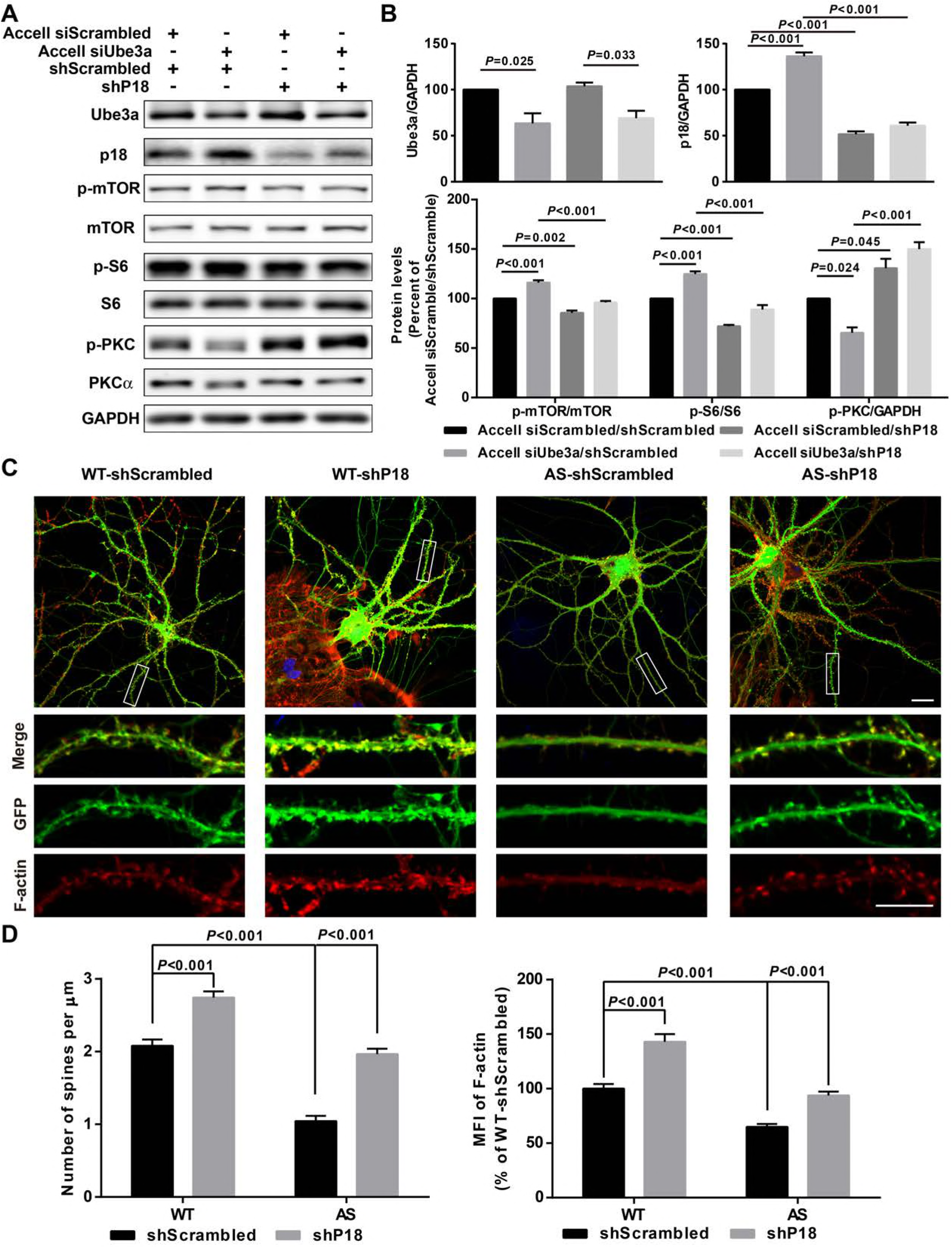
P18 mediates the effects of Ube3a on mTOR signaling, dendritic spine morphology, and actin polymerization. (A) Representative images of Western blots labeled with Ube3a, p18, p-mTOR, mTOR, p-S6, S6, p-PKC, and PKCα (GAPDH as a loading control). Protein lysates from cultured hippocampal neurons transfected with the indicated constructs were prepared for Western blot analysis. (B) Quantitative analysis of blots shown in A. N = 3, Accell siScrambled/shScrambled vs Accell siUbe3a/shScrambled, p = 0.025 (Ube3a), p < 0.001 (p18), p < 0.001 (p-mTOR), p < 0.001 (p-S6), p = 0.024 (p-PKC); Accell siScrambled/shScrambled vs Accell siScrambled/shP18, p < 0.001 (p18), p = 0.002 (p-mTOR), p < 0.001 (p-S6), p = 0.045 (p-PKC); Accell siUbe3a/shScrambled vs Accell siUbe3a/shP18, p < 0.001 (p18), p < 0.001 (p-mTOR), p < 0.001 (p-S6), p < 0.001 (p-PKC); Accell siScrambled/shP18 vs Accell siUbe3a/shP18, p = 0.033 (Ube3a); two-way ANOVA with Tukey’s post-test. (C) Representative images of F-actin (red) and GFP in cultured WT and AS hippocampal neurons (22 DIV) co-infected with GFP lentivirus and p18 shRNA or scrambled shRNA lentivirus. Scale bar, 20 μm (upper) or 10 μm (lower). (D) Quantitative analysis of images shown in C. N = 9 neurons from at least 3 independent experiments, p<0.001, two-way ANOVA with Tukey’s post-test. See also Figure 5-figure supplement 1.

We also determined whether Ube3a-mediated p18 regulation could affect dendritic spine morphology and actin polymerization. Cultured neurons from AS or WT mice were co-transfected with p18 shRNA or control shRNA lentiviruses and GFP control lentivirus, and actin polymerization was determined by staining for filamentous actin (F-actin). Confocal images of transfected neurons indicated that p18 shRNA transfection reduced p18 expression in cultured neurons from both WT and AS mice (Figure 5-figure supplement 1). Neurons from AS mice exhibited reduced dendritic spine density and actin polymerization, as compared to neurons from WT mice (Figure 5C,D). Transfection with p18 shRNA lentiviruses improved spine morphology and restored actin polymerization in neurons from AS mice, and slightly enhanced spine density and actin polymerization in neurons from WT mice (Figure 5C,D). These results indicate that deficiency in Ube3a-mediated p18 degradation contributes to spine defects and actin polymerization abnormality in neurons from AS mice.

### 6. P18 downregulation promotes LTP and improves dendritic spine morphology in AS mice

High magnification confocal images of adult hippocampal CA1 pyramidal neurons revealed that in addition to being co-localized with lysosomal markers, p18 was also localized in the vicinity of and often co-localized with PSD95 (arrowheads in Figure 6A). Quantitative analysis showed that the number of p18-immunoreactive puncta was markedly increased in AS mice (Figure 6-figure supplement 1A). Furthermore, the percentage of PSD95-stained puncta stained with p18 was also significantly increased in AS mice, as compared to WT mice (Figure 6-figure supplement 1A), although there was no significant difference in the overall number of PSD95-immunopositive puncta between AS and WT mice (Figure 6-figure supplement 1A). To determine whether increased synaptic p18 levels could contribute to impaired functional and structural synaptic plasticity in AS mice, we performed *in vivo* p18 siRNA knockdown experiments.

**Figure 6.**
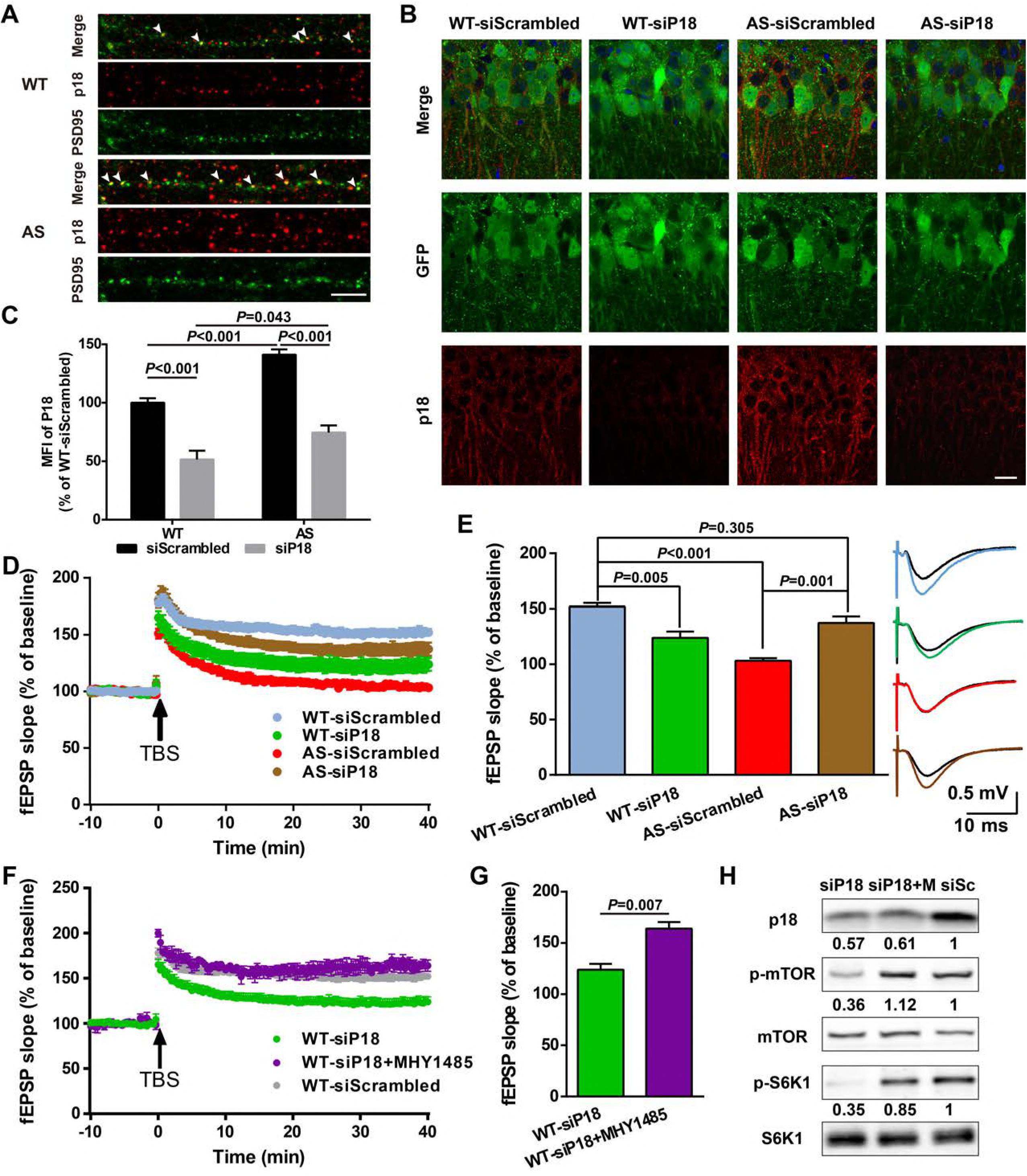
Effects of p18 downregulation in hippocampal CA1 region on LTP in WT and AS mice. (A) Representative images of dendrites of CA1 pyramidal neurons stained with anti-p18 (red) and -PSD95 (green) antibodies. Arrowheads indicate co-localized puncta. Scale bar = 10 μm. (B) Representative images of CA1 pyramidal neurons stained with anti-p18 (red) and -GFP (green) antibodies. Scale bar = 20 μm. (C) Quantitative analysis of the mean fluorescence intensity (MFI) of p18-immunoreactive puncta in GFP-positive CA1 pyramidal neurons. N = 6, p < 0.001, WT-siScrambled vs WT-siP18, p < 0.001, WT-siScrambled vs AS-siScrambled, p < 0.001, AS-siScrambled vs AS-siP18, p = 0.043, WT-siP18 vs AS-siP18, two-way ANOVA with Tukey’s post-test. (D-E) Effects of AAV siRNA-mediated p18 downregulation on LTP in WT and AS mice. (D) Slopes of fEPSPs were normalized to the average values recorded during the 10 min baseline. (E) Means ± S.E.M. of fEPSPs measured 40 min after TBS in different groups. N = 7-14 slices from 3–8 mice, p = 0.005, WT-siScrambled vs WT-siP18, p < 0.001, WT-siScrambled vs AS-siScrambled, p = 0.001, AS-siScrambled vs AS-siP18, p = 0.305, WT-siScrambled vs AS-siP18, two-way ANOVA with Tukey’s post-test. Inset shows representative traces of evoked fEPSPs before and 40 min after TBS. Scale bar 0.5 mV/10 ms. (F-G) Effects of MHY1485 treatment on LTP in p18 siRNA-injected WT mice. (F) Slopes of fEPSPs were normalized to the average values recorded during the 10 min baseline. (G) Means ± S.E.M. of fEPSPs measured 40 min after TBS in different groups. N = 3-14, p = 0.007, unpaired t-test. (H) Representative Western blots showing the relative abundance of p18, p-mTOR/mTOR, and p-S6K/S6K in lysates from control siRNA (siSc) or p18 siRNA (siP18)-transfected WT hippocampal slices. Slices were treated with or without MHY1485 (M). See also Figure 6-figure supplement 1 and 2.

AAV vectors containing p18 siRNA or scrambled siRNA (control) were bilaterally injected into the dorsal hippocampal CA1 region of WT and AS mice (Figure 6-figure supplement 1B), and LTP in hippocampal slices was analyzed 4 weeks later from these 4 experimental groups. To evaluate transfection efficiency, GFP expression, as well as p18 levels were determined in hippocampal slices by immunohistochemistry following LTP recording (Figure 6B). Only mice that exhibited significant GFP expression in the CA1 region (Figure 6-figure supplement 1C) were included in the LTP and spine analyses. Quantitative analysis showed that p18 expression was significantly increased in AS mice, as compared to WT mice, and this increase was reversed by p18 siRNA transfection (Figure 6C). P18 siRNA transfection also significantly reduced p18 expression in WT mice (Figure 6C). Levels of p18, p-mTOR/mTOR, p-S6/S6, and PKCα in CA1 regions were also determined using Western blots; p18 knockdown reduced mTORC1 activity and increased mTORC2 activity in both WT and AS mice (Figure 6-figure supplement 1D,E).

Baseline synaptic responses, including input/output curves (I/O curves) and paired-pulse facilitation, were not altered by control siRNA or p18 siRNA in either AS or WT mice (Figure 6-figure supplement 2). LTP was induced by applying theta-burst stimulation (TBS) to Schaffer collaterals in the CA1 region, as previously described (Baudry et al., 2012; Sun et al., 2015b). TBS elicited typical LTP in field CA1 of hippocampal slices from control siRNA-injected WT mice, whereas it only elicited a transient facilitation in slices from control siRNA-injected AS mice (Figure 6D,E), a result similar to that found in slices from untreated AS mice (Baudry et al., 2012; Sun et al., 2015b). In contrast, bilateral CA1 injection of p18 siRNA improved TBS-elicited LTP in hippocampal slices from AS mice (Figure 6D,E), while it reduced TBS-induced LTP in slices from WT mice (Figure 6D,E). To determine whether reduced LTP in p18 siRNA WT group was due to reduced mTORC1 activation because of “below normal” p18 levels, we used an mTOR activator MHY1485. Pre-incubation of hippocampal slices with MHY1485 (2 μM) for 60 min reestablished TBS-elicited LTP to levels identical to those in control siRNA-injected WT mice (Figure 6F,G). Levels of p18, p-mTOR/mTOR, and p-S6K1/S6K1 in AAV infected regions were determined using Western blots; p18 knockdown-induced reduction of mTORC1 activity in WT mice was reversed by acute treatment with MHY1485 (Figure 6H).

We then performed Golgi staining in hippocampal CA1 region of WT and AS mice injected with p18 siRNA or control siRNA. As previously reported (Dindot et al., 2008; Sun et al., 2016), spine density was lower with a higher proportion of immature spines (thin or filopodia) in AS mice, as compared with WT mice (Figure 7A,B and Figure 7-figure supplement 1). P18 siRNAs treatment normalized the number and improved the morphology of dendritic spines of hippocampal pyramidal neurons in AS mice, while it increased the number and proportion of immature dendritic spines in WT mice (Figure 7A,B and Figure 7-figure supplement 1). Changes in mature spine number were well related with changes observed in the LTP experiments (Figure 7B). As an independent means of assessing the effects of the changes in spine morphology on synaptic function, we recorded miniature excitatory postsynaptic currents (mEPSCs) from WT and AS hippocampal pyramidal neurons in acute slice preparations. We observed a reduction in the frequency of mEPSCs with no corresponding change in amplitude in AS neurons, as compared to WT neurons (Figure 7C,D).

**Figure 7.**
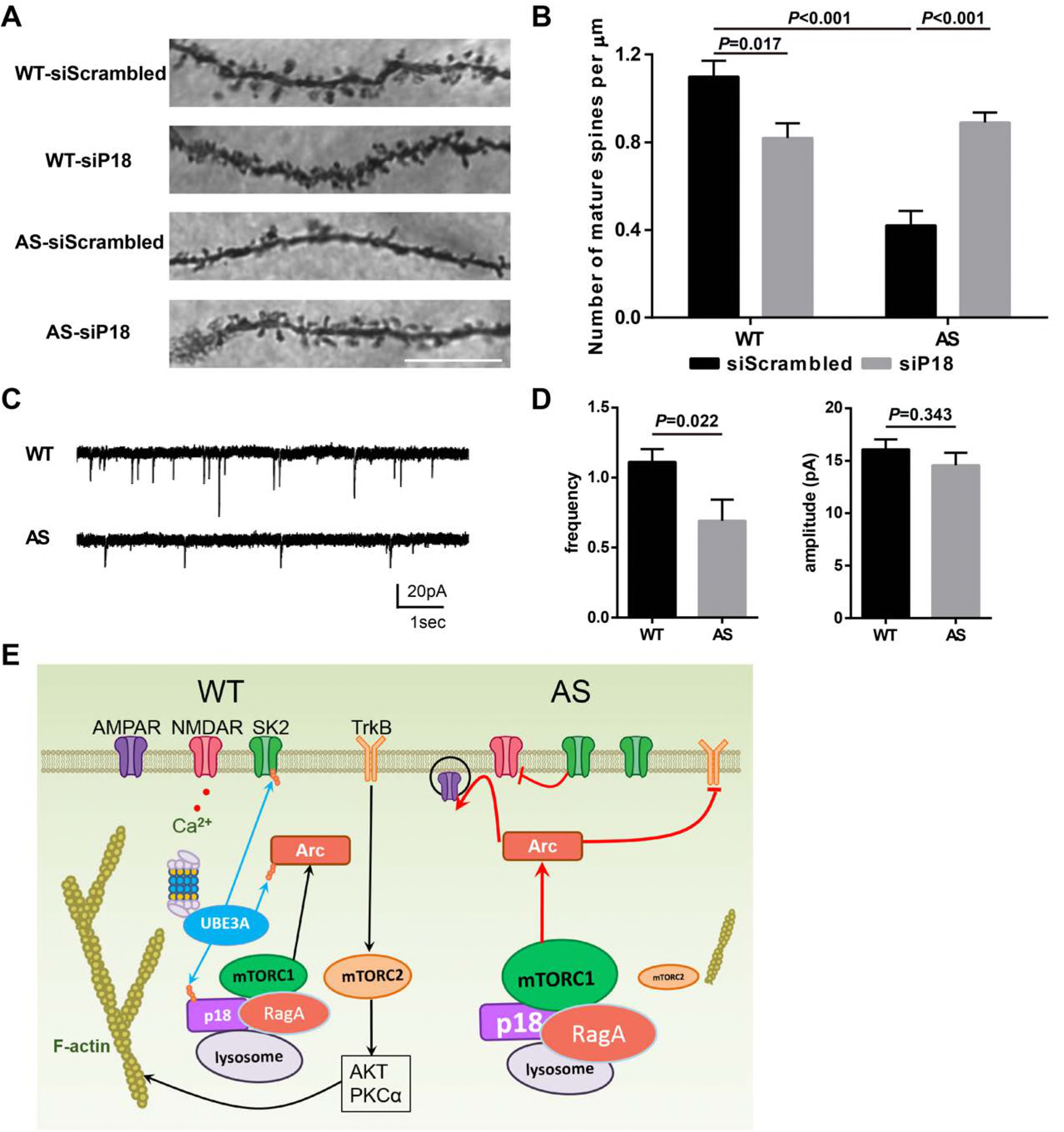
Effects of Ube3a deficiency and p18 downregulation in hippocampal CA1 region on dendritic spine morphology and mEPSCs. (A) Representative light micrograph images from Golgi-impregnated CA1 pyramidal neurons. Scale bar = 10 μm. (B) Quantitative analysis of mature dendritic spine (multi-head, mushroom, and stubby spines) density shown in A (means ± SEM from 10 slices). p = 0.017, WT-siScrambled vs WT-siP18, p < 0.001, WT-siScrambled vs AS-siScrambled, p < 0.001, AS-siScrambled vs AS-siP18, two-way ANOVA with Tukey’s post-test. (C) Representative mEPSC traces recorded in hippocampal neurons from WT and AS slices. Scale bar, 20 pA/1 s. (D) Quantification of mEPSC frequency (p = 0.022) and amplitude (p = 0.343) from WT (n = 12) and AS (n = 7) mice. Student’s t test. (E) Model for Ube3a-mediated regulation of synaptic plasticity (see text for details). See also Figure 7-figure supplement 1.

## Discussion

The last decade has witnessed a rapid growth in our knowledge of amino acid-mediated regulation of mTORC1 signaling, including the identification of a lysosome-based platform for its activation. Although regulation of mTORC1 by the TSC complex-Rheb axis is well documented in brain, its regulation by a lysosome-anchored platform has rarely been studied. We showed here for the first time that in hippocampal neurons, p18 is essential for lysosomal localization of other Ragulator members and Rag GTPases, as p18 knockdown markedly reduced the lysosomal localization of these proteins, which is in agreement with what has been reported in other cell types (Nada et al., 2009; Sancak et al., 2010). In this regard, a recent crystal structure study revealed that p18 forms a “ribbon” around the other 4 members of the Ragulator to stabilize the complex and provide contact points for Rag GTPases (de Araujo et al., 2017). We further showed that lysosomal localization of the Ragulator-Rag complex is essential for mTORC1 activation in hippocampal neurons, as previously shown in other cell types. As mentioned earlier, little is known regarding the regulation of p18 levels in any cell types. Our results provide several lines of evidence indicating that Ube3a is an E3 ligase for p18, and that Ube3a-mediated p18 ubiquitination leads to its degradation by proteasomes. Of note, p18 myristoylation and lysosomal localization of p18 were required for its ubiquitination, suggesting that Ube3a specifically targets lysosomal-localized p18, which is closely associated with mTORC1 activity regulation.

We next showed that Ube3a-mediated regulation of p18 levels is critical for mTOR signaling in hippocampal neurons, as Ube3a deficiency-induced increase in p18 levels facilitated lysosomal recruitment of Ragulator-Rag complex, leading to mTORC1 overactivation. Our results also support the idea that the Ragulator functions not only as a platform, but also as a Rag GTPase GEF, to facilitate mTORC1 activation (Bar-Peled et al., 2012). Furthermore, p18 knockdown reversed Ube3a deficiency-induced increase in mTORC1 activation and decrease in mTORC2 activation observed in AS mice. Collectively, our results indicate that Ube3a-mediated p18 ubiquitination and degradation are critical for maintaining an optimal level of lysosome-anchored Ragulator-Rag complex and a balanced mTORC1/mTORC2 activation. Our *in vivo* experiments further showed that p18 knockdown, in parallel with “normalizing” mTORC1/mTORC2 activation, restored TBS-induced LTP and improved dendritic spine morphology in hippocampus of AS mice. In support of the idea that lysosomal localized p18 contributes to synaptic plasticity regulation, we showed that in apical dendrites of hippocampal CA1 pyramidal neurons, p18 was in the vicinity of or co-localized with PSD95-labeled synapses. Furthermore, it has recently been shown that in cultured neurons lysosomes can be recruited into dendritic spines in an activity-dependent manner (Goo et al., 2017). The synaptic localization of p18 could result in the rapid assembly or disassembly of various platforms required for activation of either MAPK/ERK or mTOR complexes, which are critical for functional and structural synaptic plasticity. These lysosome-based regulatory platforms could also link local protein synthesis with local protein degradation, which would enable efficient synaptic remodeling.

Several mechanisms could account for p18 knockdown-induced LTP restoration in AS mice (see schematic in Figure 7E). First, several groups, including our own, have shown that in AS mice there is an increased expression of the immediate-early gene Arc (activity-regulated cytoskeleton-associated protein) (Cao et al., 2013; Greer et al., 2010; Sun et al., 2016). Arc has been shown to induce AMPAR endocytosis, which is an important factor in LTD (Chowdhury et al., 2006; Rial Verde et al., 2006). Related to this, both NMDAR-dependent and mGluR-dependent LTD in hippocampus are enhanced in AS mice (Pignatelli et al., 2014; Sun et al., 2015b). In AS mice, increased Arc levels were shown to impair TrkB-PSD95 signaling (Cao et al., 2013). Therefore, p18 knockdown could improve LTP by inhibiting the mTORC1-S6K1 pathway, resulting in reduced Arc protein synthesis and levels, as was the case with rapamycin treatment (Sun et al., 2016). Second, ample evidence has indicated that the dynamic properties of actin networks are crucial for synaptogenesis and synaptic plasticity, and that LTP consolidation is accompanied with increased levels of F-actin (Kramar et al., 2006; Lin et al., 2005). mTORC2 activity is also crucial for actin polymerization (He et al., 2013; Huang et al., 2013; Jacinto et al., 2004; Sun et al., 2016; Thomanetz et al., 2013). LTP impairment in AS mice was associated with reduced TBS-induced actin polymerization, as compared to WT mice, and this reduction could be ameliorated by either a positive AMPAR modulator or a SK2 channel blocker (Baudry et al., 2012; Sun et al., 2015b). Both compounds increase NMDAR activity and Ca^2+^ influx, thereby activating signaling proteins (e.g. CamKII, PKA, Rho), which facilitate F-actin formation. P18 knockdown increased mTORC2 activity, which also promotes actin polymerization, albeit through different downstream signaling pathways (activation of PKCα and Akt, etc.). Importantly, baseline synaptic transmission and paired-pulse facilitation were not altered by p18 down-regulation in both WT and AS mice, indicating that changes in synaptic plasticity resulting from p18 down-regulation are likely due to postsynaptic modifications related to processes that promote actin filament assembly during the minutes following TBS. On the other hand, our results indicated that, while there was a significant reduction in the frequency of mEPSCs in AS mice, mEPSC amplitude was not different from that measured in WT mice. Such a pattern is consistent with a loss of spines and retention of normal AMPA receptor density in the remaining, intact spine population. In accord with the latter point, we have reported that there is no change in paired-pulse facilitation or the I/O curve in AS slices (Baudry et al., 2012; Sun et al., 2015b).

P18 down-regulation slightly reduced TBS-induced LTP in slices from WT mice, and resulted in increased number and proportion of immature dendritic spines. In contrast, rapamycin treatment did not affect either LTP or spine morphology in WT mice (Sun et al., 2016). This difference is probably due to different levels of mTOR signaling alterations resulting from increased p18 levels or rapamycin treatment. Indeed, our results showed that acute activation of mTORC1 improved TBS-elicited LTP in p18 siRNA-injected WT slices. Furthermore, rapamycin treatment (at the dose we used) had no effect on the level of p-mTOR 2448 (Sun et al., 2016), while p18 silencing significantly reduced the level of p-mTOR 2448. Of note, work from the Costa-Mattioli lab has clearly indicated that a low concentration of rapamycin has no effect on LTP in WT mice, while a high concentration of rapamycin impairs LTP (Stoica et al., 2011). Collectively, these results further support the notion that optimal p18 levels and mTORC1 activation are necessary for synaptic plasticity. In light of a recent study showing that Ube3a plays an important role in experience-dependent maturation of excitatory synapses in the visual cortex (Kim et al., 2016), it is conceivable that Ube3a-mediated p18 regulation may also contribute to synaptic plasticity and circuit remodeling in other brain areas. P18 has been shown to directly interact with p27 (kip1), thereby regulating RhoA activity and actin remodeling (Hoshino et al., 2011), and autophagic activity (Zada et al., 2015). Whether these mTOR-independent p18 functions play any role in synaptic plasticity and brain development remains to be determined.

Although we propose that Ube3a-mediated regulation of p18-mTOR pathway is crucial in the pathogenesis of AS, our work by no means intends to conclude that p18 is the sole Ube3a target responsible for AS. Rather, our results indicate that the newly identified regulation of mTORC1 activation by lysosome-located p18 is present in brain and plays important roles in synaptic plasticity, and document the existence of another layer of regulation in the already complex mTOR pathway, namely the regulation of p18 levels by Ube3a. Of note, while UBE3A deficiency results in AS, UBE3A over-expression results in increased ASD risk. An increased density of dendritic spines with immature morphology has been reported in brains of ASD patients (Hutsler and Zhang, 2010; Tang et al., 2014). In our study, reducing p18 levels in WT mice resulted in similar changes in spine properties. It is therefore tempting to propose that UBE3A over-expression might induce ASD phenotypes, at least in part, by down-regulating p18 levels. It is also noteworthy that abnormal mTOR signaling has been implicated in a number of diseases. Therefore, results from our studies, shed new light on a broad range of normal brain functions, and on several neurological/neuropsychiatric disorders, including AS.

## Author Contributions

X.B. and M.B. designed research; J.S., Y.L., Y.J., J.T., X.H., and W.L. performed research and analyzed data; J.S., G.L., M.B. and X.B. wrote the paper. All authors reviewed the manuscript.

## Acknowledgments

This work was supported by grant P01NS045260-01 from NINDS (PI: Dr. C.M. Gall), and grants R01NS057128 from NINDS to MB and R15MH101703 from NIMH to XB. XB is also supported by funds from the Daljit and Elaine Sarkaria Chair. The authors declare no conflicts of interest.

## Materials and Methods

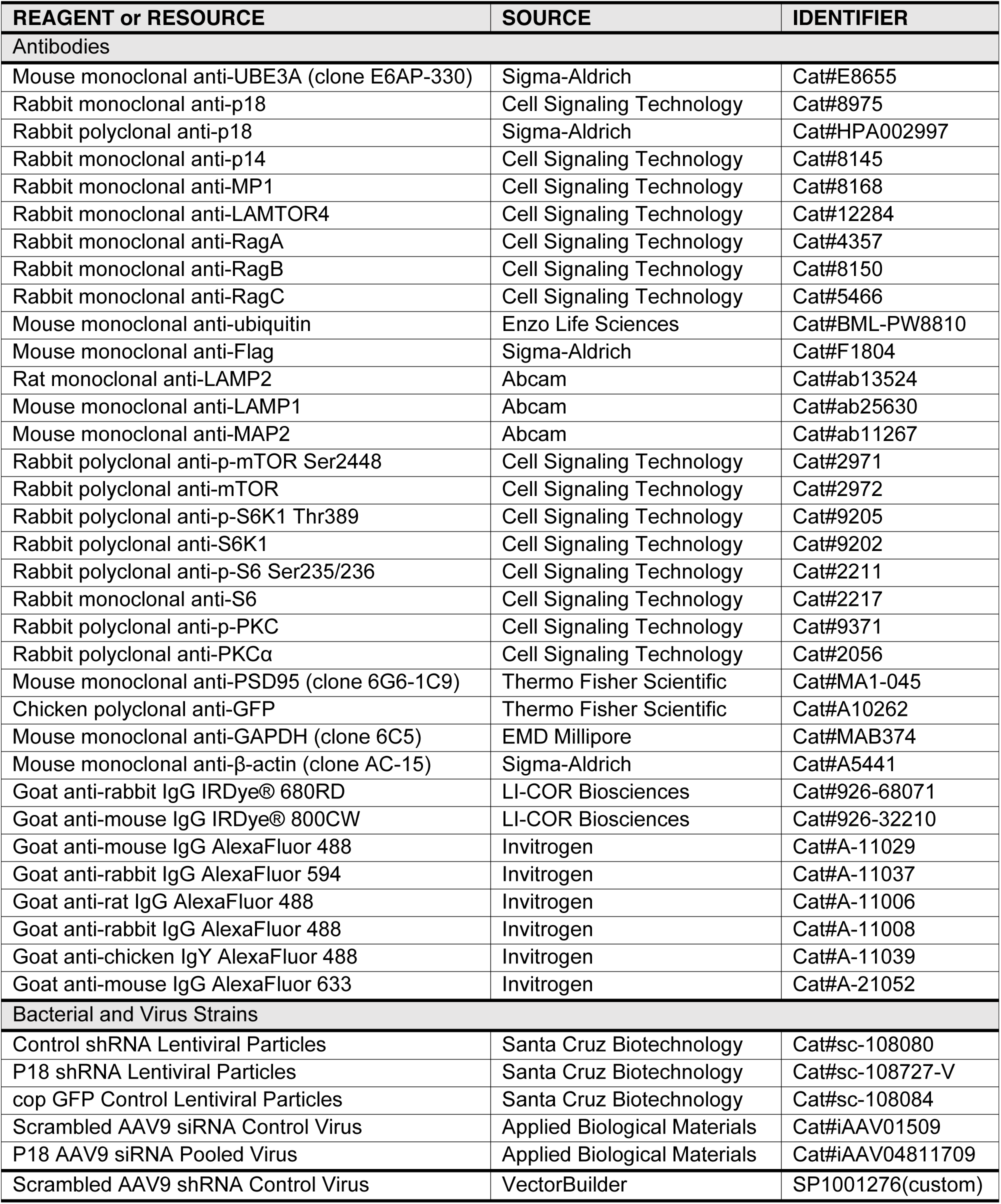
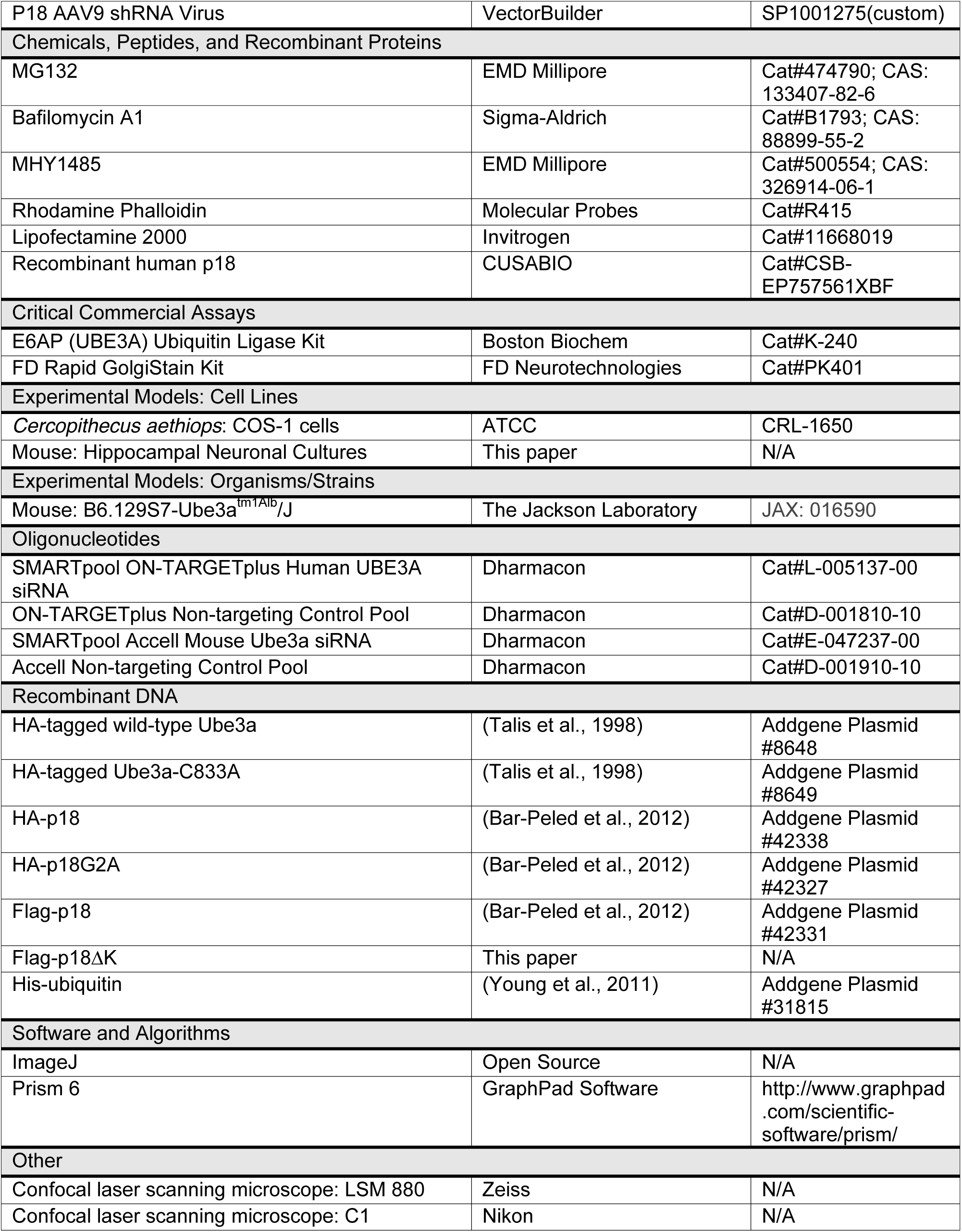
KEY RESOURCES TABLE.

## EXPERIMENTAL MODEL AND SUBJECT DETAILS

### Animals

Animal experiments were conducted in accordance with the principles and procedures of the National Institutes of Health Guide for the Care and Use of Laboratory Animals. All protocols were approved by the local Institutional Animal Care and Use Committee. Original Ube3a mutant (AS) mice were obtained from The Jackson Laboratory, strain B6.129S7-Ube3a^tm1Alb^/J, and a breeding colony was established, as previously described (Baudry et al., 2012). In all experiments male AS mice aged between 2–4 months were used. Control mice were age-matched, male, wild-type littermates. Mice, housed in groups of two to three per cage, were maintained on a 12-h light/dark cycle with food and water ad libitum.

### Hippocampal Neuronal Cultures

For neuronal culture preparations, wild-type (WT) or Ube3a^m-/p+^ female and WT male mice were used for breeding. Hippocampal neurons were prepared from E18 mouse embryos as described (Sun et al., 2015b). Briefly, hippocampi were dissected and digested with papain (2 mg/ml, Sigma) for 30 min at 37 °C. Dissociated cells were plated onto poly-D-lysine-coated 6-well plate at a density of 6-10×10^4^ cells/cm^2^ or coverslips in 24-well plate at a density of 6-10×10^3^ cells/cm^2^ in Neurobasal medium (GIBCO) supplemented with 2% B27 (GIBCO) and 2 mM glutamine and kept at 37 °C under 5% CO_2_. Half of the culture medium was replaced with fresh culture medium at DIV4 and then every 7 days. Genotyping was carried out by polymerase chain reaction (PCR) of mouse tail DNA as described previously (Baudry et al., 2012).

### Cell Lines

COS-1 cells (ATCC) were grown in DMEM supplemented with 10% (vol/vol) fetal bovine serum (FBS) (Invitrogen) and kept at 37 °C under 5% CO_2_.

## METHODS

### Transfection and Lentiviral Infection

For transient expression of constructs, COS-1 cells were transfected with the respective constructs by lipofection (Lipofectamine 2000; Invitrogen) according to the manufacturer’s instructions. Small interfering RNA (siRNA) transfections were also performed with Lipofectamine 2000. Cells were incubated with 10 or 20 nM SMARTpool siRNA duplexes against human *UBE3A*, or a scrambled duplex (Dharmacon) for 72 h before downstream analysis.

Cultured hippocampal neurons from WT mice were infected with p18 shRNA (mouse) lentivirus (sc-108727-V, Santa Cruz Biotechnology) or scrambled shRNA lentivirus (sc-108080, Santa Cruz Biotechnology), and co-transfected with Accell Ube3a siRNA (GE Dharmacon) or Accell Non-targeting siRNA (GE Dharmacon) at DIV 4, and 24 h after infection, 2/3 medium was replaced with fresh medium. Cultured neurons were used 3 days after infection.

Cultured WT and AS hippocampal neurons were infected with p18 shRNA (mouse) lentivirus (sc-108727-V, Santa Cruz Biotechnology) or scrambled shRNA lentivirus (sc-108080), together with copGFP control lentivirus (as an infection marker, sc-108084) at DIV 14, and 24 h after infection, 2/3 medium was replaced with fresh medium. Neurons were analyzed 8 days after infection.

Cultured WT and AS hippocampal neurons were infected with p18 shRNA (mouse) AAV (custom, VectorBuilder) or scrambled shRNA AAV (custom, VectorBuilder) at DIV 7, and 24 h after infection, 2/3 medium was replaced with fresh medium. Neurons were analyzed 14 days after infection.

### Antibodies

The following primary antibodies were used: UBE3A (Sigma, E8655), p18 (CST, 8975), p18 (Sigma, HPA002997), p14 (CST, 8145), MP1 (CST, 8168), LAMTOR4 (CST, 12284), RagA (CST, 4357), RagB (CST, 8150), RagC (CST, 5466), ubiquitin (Ub, Enzo, BML-PW8810), Flag (Sigma, F3165), LAMP2 (Abcam, ab13524), LAMP1 (Abcam, ab25630), MAP2 (Abcam, ab11267), p-mTOR Ser2448 (CST, 2971), mTOR (CST, 2972), p-S6K1 (CST, 9205), S6K1 (CST, 9202), p-S6 Ser235/236 (CST, 2211), S6 (CST, 2217), p-PKC (CST, 9371), PKCα (CST, 2056), PSD95 (Thermo, MA1-045), GFP (Thermo, A10262), GAPDH (Millipore, MAB374), and β-actin (Sigma, A5441). All secondary antibodies for Western blots were obtained from LI-COR, and for immunofluorescence Alexa-488, -594, and -633 conjugated secondary antibodies were obtained from Invitrogen.

### DNA Constructs

Expression constructs encoding HA-tagged wild-type Ube3a, HA-tagged catalytically inactive mutant Ube3a-C833A (substitution of Cys-833 by Ala), HA-p18, HA-p18G2A (substitution of Gly-2 by Ala), Flag-p18, and His-ubiquitin were obtained from Addgene. Constructs with K-R mutations (Flag-p18∆K) were generated from Flag-p18 by arginine substitution of all lysine residues.

### P2/S2 Fractionation and Western Blot Analysis

P2/S2 fractionation were performed according to published protocols (Sun et al., 2015a). Briefly, frozen hippocampus tissue was homogenized in ice-cold HEPES-buffered sucrose solution (0.32 M sucrose, 4 mM HEPES, pH 7.4) with protease inhibitors. Homogenates were centrifuged at 900 g for 10 min to remove large debris (P1). The supernatant (S1) was then centrifuged at 11,000 g for 20 min to obtain crude membrane (P2) and cytosolic (S2) fractions. P2 pellets were sonicated in RIPA buffer (10 mM Tris, pH 8, 140 mM NaCl, 1 mM EDTA, 0.5 mM EGTA, 1% NP-40, 0.5% sodium deoxycholate, and 0.1% SDS). For whole homogenates, tissue was homogenized in RIPA buffer. Protein concentrations were determined with a BCA protein assay kit (Pierce).

Western blots were performed according to published protocols (Sun et al., 2015b). Briefly, samples were separated by SDS-PAGE and transferred onto a PVDF membrane (Millipore). After blocking with 3% BSA for 1 h, membranes were incubated with specific antibodies overnight at 4 °C followed by incubation with secondary antibodies (IRDye secondary antibodies) for 2 h at room temperature. Antibody binding was detected with the Odyssey^®^ family of imaging systems.

### Immunoprecipitation and Denaturing Immunoprecipitation

For immunoprecipitation, all procedures were carried out at 4 °C. COS-1 cells transfected with the indicated cDNAs or cultured hippocampal neurons were lysed with lysis buffer (Tris-HCl 25 mM pH 7.4, NaCl 150 mM, 1 mM EDTA, 1% NP-40, 5% glycerol and a protease inhibitor cocktail). After a brief centrifugation to remove insoluble material, the supernatant was precleared with an aliquot of agarose beads. For immunoprecipitation of Flag-p18 or Flag-p18∆K in COS-1 cells, extracts were incubated overnight with anti-Flag agarose beads, washed with lysis buffer, followed by elution of bound proteins by heating at 100 °C for 10 min in SDS-PAGE sample buffer. For immunoprecipitation of p18 in hippocampal neurons, extracts were incubated with anti-p18 antibodies overnight and immunoprecipitates were collected with protein A/G Agarose. For immunoprecipitation of RagA in mouse hippocampus, rabbit anti-RagA antibodies were incubated with hippocampal lysates and precipitated with Protein A/G-conjugated beads. Inputs and precipitates were resolved by SDS-PAGE and analyzed by Western blotting. All studies were performed in 3–5 independent experiments.

For immunoprecipitation of ubiquitin from hippocampal crude membrane fractions under denaturing conditions, P2 pellets were resuspended and heated in denaturing lysis buffer (1 % SDS, 50 mM Tris, pH 7.4, 5 mM EDTA, 10 mM DTT, 1 mM PMSF, 2 μg/ml leupeptin, 15 U/ml DNase I) and diluted in 9 volumes of ice-cold non-denaturing lysis buffer (1 % Triton X-100, 50 mM Tris, pH 7.4, 300 mM NaCl, 5 mM EDTA, 10 mM iodoacetamide, 1 mM PMSF, 2 μg/ml leupeptin). Lysates were centrifuged at 16,000 g for 30 min at 4 °C and cleared with protein A/ G Agarose beads. Pre-cleared lysates were then incubated with anti-ubiquitin antibodies coupled to protein A/G Agarose beads overnight at 4°C, followed by four washes with ice-cold wash buffer (0.1 % Triton X-100, 50 mM Tris, pH 7.4, 300 mM NaCl, 5 mM EDTA) and elution in 2 x SDS sample buffer. Immunoprecipitated proteins were resolved by SDS-PAGE followed by Western blot analysis with specific antibodies against p18 and ubiquitin. At least three independent experiments were performed.

### *In Vitro* Ubiquitin Assay

His-p18 proteins were purchased from CUSABIO (Wuhan, China). For *in vitro* ubiquitination experiments, we used the E6AP (UBE3A) Ubiquitin Ligase Kit (Boston Biochem), following the manufacturer’s instruction. Briefly, purified His-p18 proteins were incubated for 90 min at 37 °C under constant shaking with E1 enzyme, E2 enzyme (UBE2L3), His_6_-E6AP, ubiquitin, Mg^2+^-ATP, and Reaction Buffer. The reaction was terminated by the addition of SDS sample buffer, and samples were boiled, and proteins separated with 14% SDS-PAGE. Blots were probed with p18, ubiquitin, and His antibodies. At least three independent experiments were performed.

### His-ubiquitin Pull-down Assay

COS-1 cells in 60-mm dishes were transfected with 2.5 μg His-ubiquitin, 2.5 μg HA-p18 or HA-p18G2A, and 5 μg HA-Ube3a or HA-Ube3a C833A constructs in the indicated combinations. Ube3a siRNA-treated COS-1 cells were transfected with 2.5 μg Flag-p18 or p18∆K and 2.5 μg His-ubiquitin 48 h after siRNA treatment. Twenty-four hours after transfection, cells were lysed, and His-ubiquitin-conjugated proteins were purified as described (Sun et al., 2015b). Briefly, cells were harvested in 1 ml of ice-cold phosphate-buffered saline, and the cell suspension was divided into two parts; 100 μl were lysed using 1 x SDS-PAGE sample loading buffer containing 10 % DTT, and 900 μl were lysed in Buffer A (6 M guanidine HCl, 0.1 M Na_2_HPO_4_/NaH_2_PO_4_, 0.5 M NaCl, 10 mM imidazole, 0.1 % Nonidet P-40, and 5 % glycerol, pH 8.0) and sonicated. The guanidine lysates were incubated with 30 μl of equilibrated Talon resin at 4 °C for 4 h to bind His-tagged ubiquitinated proteins. Beads were then washed one time with Buffer A, followed by four washes with Buffer B (8 M urea, 0.1 M Na_2_HPO_4_/NaH_2_PO_4_, 0.5 M NaCl, 20 mM imidazole, 0.1 % Nonidet P-40, and 5 % glycerol, pH 8.0). The protein conjugates were eluted in 30 μL of 2 X laemmli/imidazole (200 mM imidazole) and boiled. Eluates were analyzed by Western blotting using either p18 or ubiquitin antibody. At least three independent experiments were performed.

### Acute Hippocampal Slice Preparation

Adult male mice (2–4-month-old) were anesthetized with gaseous isoflurane and decapitated. Brains were quickly removed and transferred to oxygenated, ice-cold cutting medium (in mM): 124 NaCl, 26 NaHCO_3_, 10 glucose, 3 KCl, 1.25 KH_2_PO_4_, 5 MgSO_4_, and 3.4 CaCl_2_. Hippocampal transversal slices (400 μm-thick) were prepared using a McIlwain-type tissue chopper and transferred to i) an interface recording chamber and exposed to a warm, humidified atmosphere of 95 % O_2_/5 % CO_2_ and continuously perfused with oxygenated and preheated (33 ± 0.5 °C) artificial cerebrospinal fluid (aCSF) (in mM) [110 NaCl, 5 KCl, 2.5 CaCl_2_, 1.5 MgSO_4_, 1.24 KH_2_PO_4_, 10 D-glucose, 27.4 NaHCO_3_], perfused at 1.4 ml/min (electrophysiology); or ii) a recovery chamber with a modified aCSF medium, containing (in mM): 124 NaCl, 2.5 KCl, 2.5 CaCl_2_, 1.5 MgSO_4_, 1.25 NaH_2_PO_4_, 24 NaHCO_3_, 10 D-glucose, and saturated with 95 % O_2_ /5 % CO_2_ for 1 h at 37 °C (biochemical assays).

### Electrophysiology

After 1.5 h incubation at 33 ± 0.5 °C in the recording chamber, a single glass pipette filled with 2 M NaCl was used to record field EPSPs (fEPSPs) elicited by stimulation of the Schaffer collateral pathway with twisted nichrome wires (single bare wire diameter, 50 μm) placed in CA1 stratum radiatum. Stimulation pulses were generated using a Multichannel Systems Model STG4002 Stimulator (Reutlingen, Germany). Responses were recorded through a differential amplifier (DAM 50, World Precision Instruments, USA) with a 10-kHz high-pass and 0.1-Hz low-pass filter. Before each experiment, the input/output (I/O) relation was examined by varying the intensity of the stimulation. Paired-pulse facilitation was tested at 20–300 ms interval. Long-term potentiation (LTP) was induced using theta burst stimulation (10 bursts at 5 Hz, each burst consisting of 4 pulses at 100 Hz, with a pulse duration of 0.2 ms). For LTP and paired-pulse facilitation experiments, the stimulation intensity was regulated to a current which elicited a 40 % of maximal response. Data were collected and digitized by Clampex, and the slope of fEPSP was analyzed. MHY1485 (2 μM) was applied to slices for 60 min before theta-burst stimulation (TBS). Some of the slices were processed for Western blots. All data are expressed as means ± SEM, and statistical significance of differences between means was calculated with appropriate statistical tests as indicated in figure legends.

Whole-cell patch-clamp recording was performed as previously described (Vogel-Ciernia et al., 2013). Briefly, hippocampal slices were prepared on the horizontal plane at a thickness of 370 μm from 2- to 4-month-old male mice with a Leica vibrating tissue slicer (Model: VT1000S). Slices were placed in a submerged recording chamber and continuously perfused at 2–3 mL/min with oxygenated (95% O_2_/5% CO_2_) at 32 °C. Whole-cell recordings (Axopatch 200A amplifier: Molecular Devices) were made with 4–7 MΩ recording pipettes filled with a solution containing (in mM): 130 CsMeSO4, 10 CsCl, 8 NaCl, 10 HEPES, 0.2 EGTA, 5 QX-314, 2 Mg-ATP, 0.3 Na-GTP. Osmolarity was adjusted to 290–295 mOsm and pH 7.4. Spontaneous mEPSCs were recorded at a holding potential of -70 mV in the presence of tetrodotoxin (1 μM) and picrotoxin (50 μM). Data were filtered at 2 kHz, digitized at 1–5 kHz, stored on a computer, and analyzed off-line using Mini Analysis Program (Synaptosoft), Origin (OriginLab) and pCLAMP 7 (Molecular Devices) software. Statistical significance was determined by pooling events from cells of the same genotype and running a Student’s t test on the pooled data. P < 0.05 was considered statistically significant.

### Immunofluorescence

Cultured hippocampal neurons were fixed in 2% paraformaldehyde (PFA)/10% sucrose for 15 min at 37 °C, transferred to 0.05% Triton X-100/PBS for 5 min at 4 °C, and then 0.02% Tween-20/PBS for 2 min at 4 °C. Coverslips were washed twice with ice cold PBS and incubated 1 h in 3% BSA/PBS at room temperature. For staining of F-actin, Rhodamine-Phalloidin was incubated in 1% BSA/PBS overnight at 4 °C. For staining of p18, LAMTOR4, RagA, LAMP2, and MAP2, cells were incubated with rabbit anti-p18 (1:200, Sigma), rabbit anti-LAMTOR4 (1:500, CST), rabbit anti-RagA (1:100, CST), rat anti-LAMP2 (1:200, Abcam), mouse anti-MAP2 (1:500, Abcam) respectively in 3% BSA/PBS overnight at 4 °C. Coverslips were then washed twice with ice cold PBS for 10 min each and then incubated with secondary antibodies (Alexa Fluor-594 anti-rabbit, 1:200; Alexa Fluor-594 anti-rat, 1:200; and Alexa Fluor-633 anti-mouse, 1:200) for 2 h at room temperature. Coverslips were then washed four times with ice cold PBS for 10 min each, and mounted on glass slides using VECTASHIELD mounting medium with DAPI (Vector Laboratories). Images were acquired using a Zeiss LSM 880 confocal laser-scanning microscope. The staining was visualized in GFP-expressed neurons.

Hippocampal slices were collected 40 min after TBS and fixed in 4% PFA for 1 h and cryoprotected in 30% sucrose for 1 h at 4 °C, and sectioned on a freezing microtome at 20 µm. Sections were blocked in 0.1 M PBS containing 5% goat serum and 0.3% Triton X-100, and then incubated in primary antibody mixture including chicken anti-GFP (1:500) and rabbit anti-p18 (1:200, Sigma) in 0.1 M PBS containing 1% BSA and 0.3% Triton X-100 overnight at 4 °C. Sections were washed 3 times (10 min each) in PBS and incubated in Alexa Fluor 488 goat anti-chicken IgG and Alexa Fluor 594 goat anti-rabbit IgG for 2 h at room temperature. All images were taken in CA1 stratum radiatum between the stimulating and recording electrodes. The threshold for the GFP fluorescence was set to make sure that the control slices from naive mice or mice with AAV infection but without GFP reporter were considered GFP-negative.

For immunofluorescence with brain tissue section, deeply anesthetized animals were perfused and brains were post-fixed in 4% PFA overnight followed by sequential immersion in 15% and 30% sucrose for cryoprotection. Brains were then sectioned (20 µm) and stained as described above. The following primary antibodies were used: p18 (1:200, Sigma), LAMTOR4 (1:500, CST), RagA (1:100, CST), p14 (1:100, CST), MP1 (1:100, CST), RagB (1:100, CST) mTOR (1:100, CST), p-mTOR (1:100, CST), LAMP2 (1:200, Abcam), and PSD95 (1:200, Thermo). The hippocampal CA1 pyramidal cell soma and apical dendrites were randomly selected for colocalization analysis by Manders’ coefficients. The apical dendrites in hippocampal CA1 stratum radiatum were also randomly selected for puncta analysis. The puncta number of p18/PSD95 was quantified and the percentage of p18 and PSD95 dually stained synapses was also analyzed.

### Intrahippocampal AAV Injection

Stereotaxic AAV injection into CA1 region of the hippocampus was performed in 8-week-old mice. Animals were allocated into the experimental/control groups in a randomized manner. Under isoflurane anesthesia, AAV p18 siRNA or AAV scrambled siRNA constructs in 2 µl solution were injected bilaterally into CA1 regions at two sites: 1.94 mm posterior to bregma, 1.4 mm lateral to the midline and 1.35 mm below the dura and 2.2 mm posterior to bregma, 1.8 mm lateral to the midline and 1.5 mm below the dura. The solution was slowly injected over 30 min and the needle was left in place for an additional 10 min. The needle was then slowly withdrawn and the incision closed. AAV-injected mice were used for experiments after four weeks, a period determined in pilot studies to be necessary for sufficient expression of viral mediated gene expression.

### Image Analysis and Quantification

Images were acquired using a Nikon C1 or a Zeiss LSM 880 with Airyscan confocal laser-scanning microscope with a 60 X objective. Images for all groups in a particular experiment were obtained using identical acquisition parameters and analyzed using ImageJ software (NIH). All immunostaining studies were performed in 3–5 independent experiments. In all cases the experimenter was blind regarding the identity of the transfected constructs and the genotypes during acquisition and analysis.

### Dendritic Spine Analysis

Four weeks after AAV injection, mice were deeply anesthetized using gaseous isoflurane and then decapitated. The brain was rapidly removed and Golgi impregnation was performed according to our published protocol (Sun et al., 2016) and outlined in the FD Rapid GolgiStain Kit (FD Neurotechnologies, Ellicott, MD). The number of spines located on randomly selected dendritic branches was counted manually by an investigator blind to genotype and injection. Spine density was calculated by dividing the number of spines on a segment by the length of the segment and was expressed as the number of spines per μm of dendrite. Spine types were determined on the basis of the ratio of the width of the spine head to the length of the spine neck and classified as previously described (Risher et al., 2015). Five to seven dendritic branches between 10 and 20 µm in length were analyzed and averaged to provide a section mean.

### Statistical Analysis

Error bars indicate standard error of the mean. To compute p values, unpaired Student’s t test, and one- or two-way ANOVA with Tukey’s post-test were used (GraphPad Prism 6), as indicated in figure legends. The level of statistical significance was set at p < 0.05.

**Figure 1-figure supplement 1.**
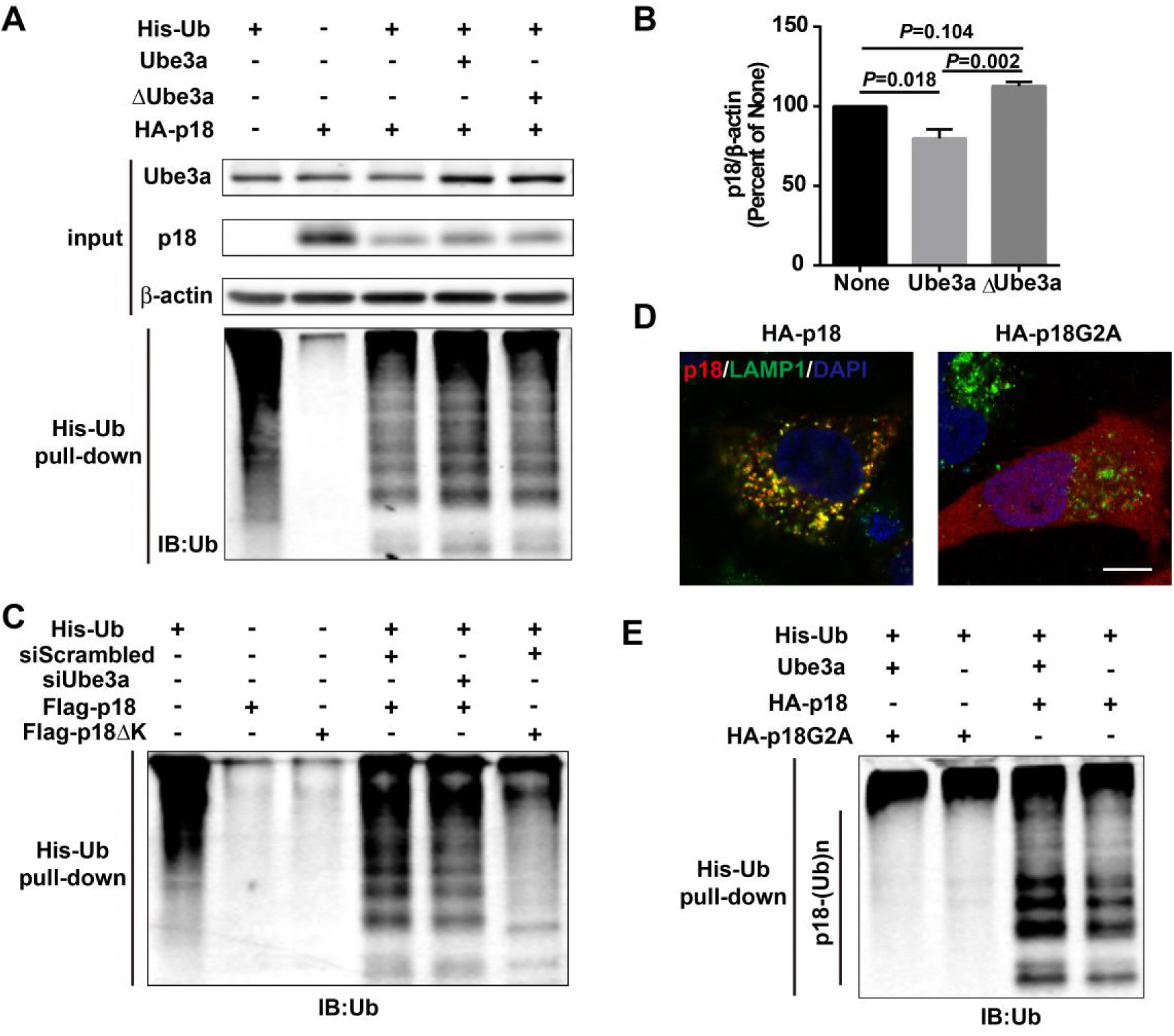
His-ubiquitin pull down analyses of p18 ubiquitination. Related to Figure 1. (A) His-ubiquitin pull down assay performed following overexpression of Ube3a or ∆∆Ube3a. Upper panel: Levels of input proteins were evaluated by Western blot probed with Ube3a, p18, and β-actin antibodies. Lower panel: Levels of ubiquitin were determined by Western blot analysis. This image is paired with Figure 1E. (B) Quantitative analysis of blots in Figure 1F (means ± SEM, p = 0.018 None vs Ube3a, p = 0.002 Ube3a vs ∆Ube3a, p = 0.104 None vs ∆Ube3a, n = 3, one-way ANOVA with Tukey’s post hoc analysis). (C) His-ubiquitin pull down assay performed following Ube3a siRNA treatment. Levels of ubiquitin were determined by Western blot analysis. This image is paired with Figure 1G. (D) Localization of wild-type p18 and p18G2A mutant. COS-1 cells expressing p18 or p18G2A were stained with anti-p18 antibody (red) and anti-LAMP1 antibody (green). Scale bar = 10 μm. (E) His-ubiquitin pull down assay performed using HA-p18 or HA-p18G2A. Levels of ubiquitin were determined by Western blot analysis. This image is paired with Figure 1H.

**Figure 2-figure supplement 1.**
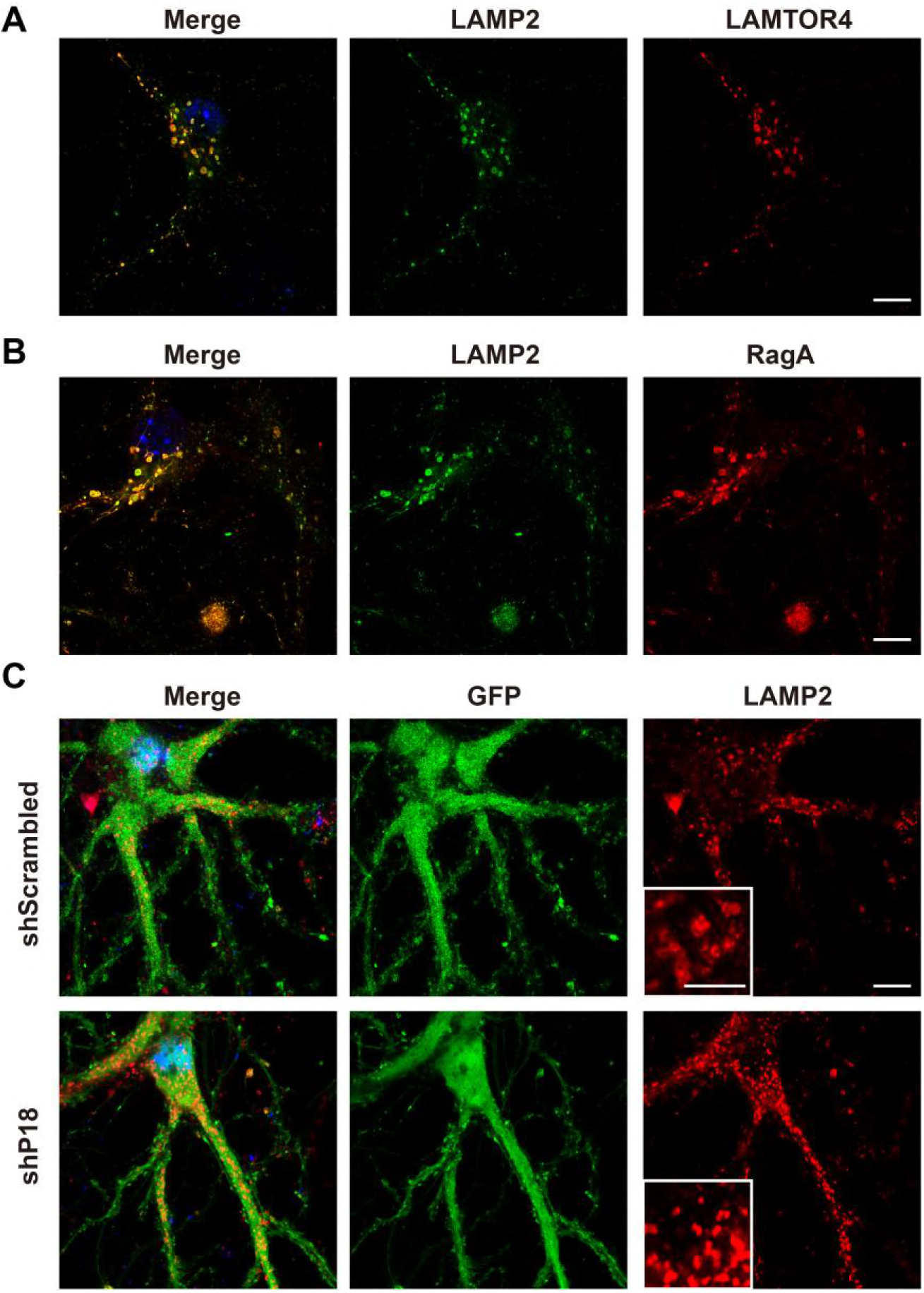
Localization of LAMTOR4 and RagA, and effects of p18 knockdown on LAMP2 in hippocampal neurons. Related to Figure 2. (A) Images of cultured hippocampal neurons co-immunostained for lysosomal protein LAMP2 (green) and LAMTOR4 (red). Scale bar = 10 μm. (B) Images of hippocampal neurons co-immunostained for lysosomal protein LAMP2 (green) and RagA (red). Scale bar =10 μm. (C) Images of hippocampal neurons stained for LAMP2 (red). Neurons were infected with shRNA AAV directed against p18 with GFP co-expression or scrambled shRNA control. Scale bar, 10 μm; inset, 5 μm.

**Figure 3-figure supplement 1.**
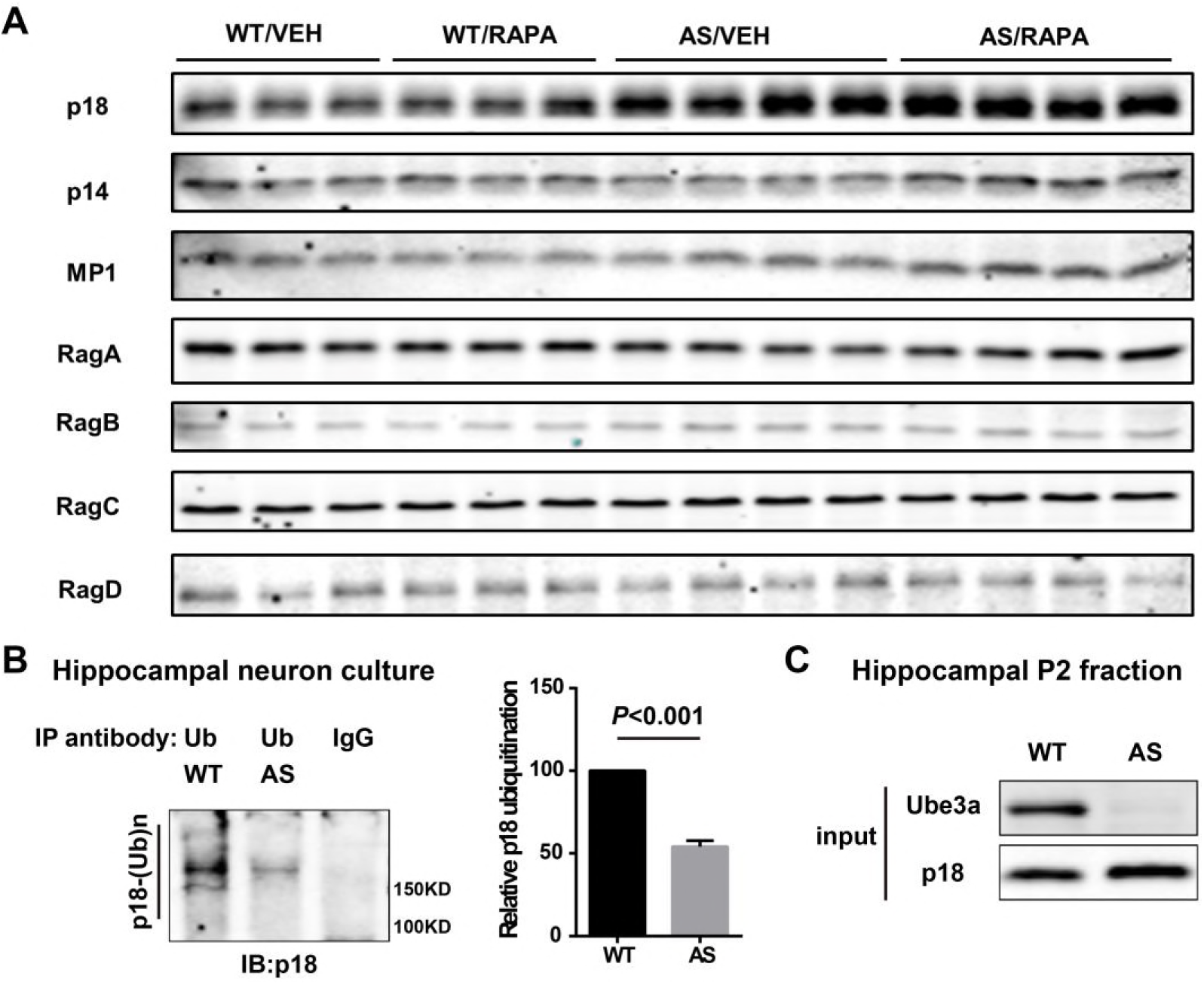
Levels of members of the Ragulator-Rag complex in WT and AS mice and *in vivo* denaturing immunoprecipitation assay of p18 ubiquitination. Related to Figure 3. (A) Representative images of Western blots labeled with p18, p14, MP1, RagA, RagB, RagC, and RagD in P2 fractions of hippocampus from vehicle (VEH)- and rapamycin (RAPA)-treated WT and AS mice. (B) Immunoprecipitation of lysates from WT and AS hippocampal neuron cultures under denaturing conditions was performed with anti-ubiquitin antibodies or control IgG and Western blots were labelled with anti-p18 antibodies. Ubiquitinated p18 proteins are indicated as “p18-(Ub)n”. Right panel: quantification of the relative abundance of ubiquitinated p18 (mean ± SEM, p < 0.001, n = 3, Student’s t test). (C) Levels of input proteins were evaluated by Western blot probed with Ube3a and p18 antibodies. This image is paired with Figure 3C.

**Figure 4-figure supplement 1.**
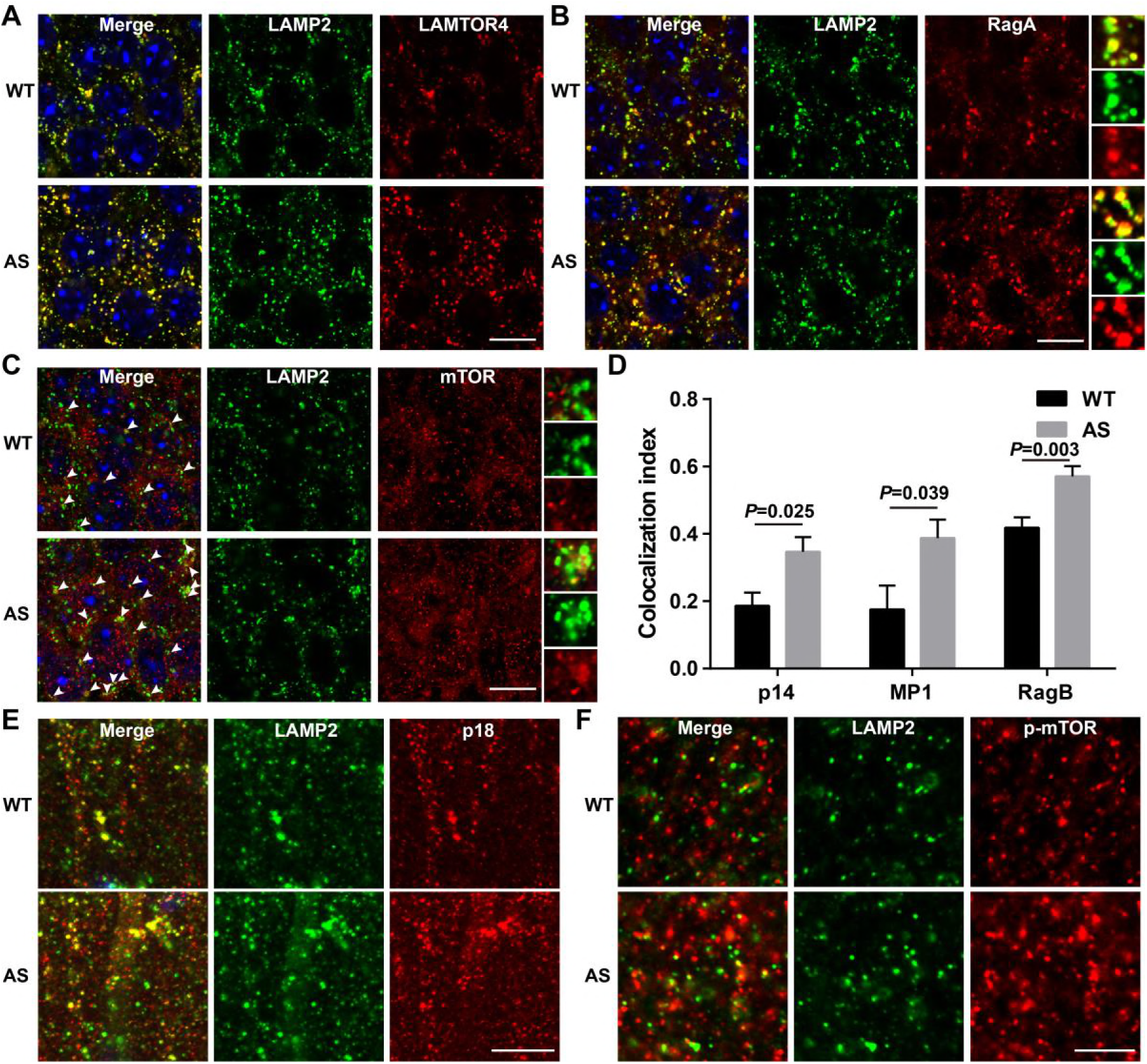
Lysosomal localization of members of the Ragulator-Rag complex and mTOR/p-mTOR in hippocampus of WT and AS mice. Related to Figure 4. (A) Co-localization of LAMTOR4 (red) with LAMP2 (green) in cell bodies of CA1 pyramidal neurons from WT and AS mice. Scale bar = 10 μm. (B) Co-localization of RagA (red) with LAMP2 (green) in cell bodies of CA1 pyramidal neurons from WT and AS mice. Scale bar = 10 μm. (C) Co-localization of mTOR (red) with LAMP2 (green) in cell bodies of CA1 pyramidal neurons from WT and AS mice. Arrowheads indicate co-localized puncta. Scale bar = 10 μm. In B and C, insets show selected fields that were magnified ten times. (D) Quantification of p14-LAMP2 (n = 5, p = 0.025), MP1-LAMP2 (n = 6, p = 0.039), and RagB-LAMP2 (n = 11 for WT, 9 for AS, p = 0.003) colocalization in cell bodies of CA1 pyramidal neurons from WT and AS mice. Unpaired t-test. (E) Representative images of apical dendrites of CA1 pyramidal neurons stained with anti-p18 (red) and -LAMP2 (green) antibodies. Scale bar = 10 μm. (F) Representative images of apical dendrites of CA1 pyramidal neurons stained with anti-p-mTOR (red) and - LAMP2 (green) antibodies. Scale bar = 5 μm.

**Figure 5-figure supplement 1.**
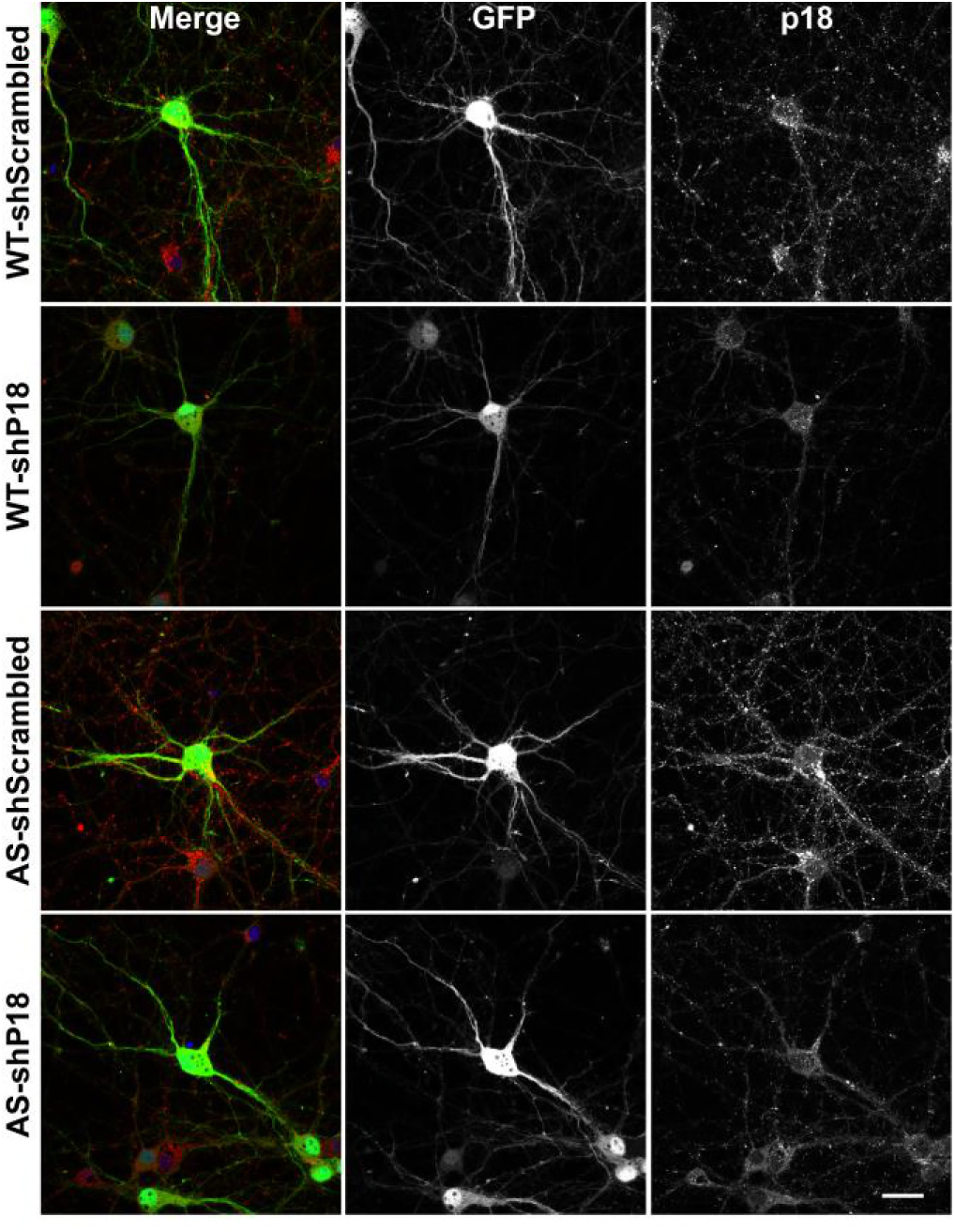
Effects of p18 knockdown in WT and AS hippocampal neurons. Related to Figure 5. Representative images of p18 (red) and GFP in WT and AS hippocampal neurons co-infected with copGFP lentivirus and p18 shRNA or scrambled shRNA lentivirus. Scale bar, 30 μm.

**Figure 6-figure supplement 1.**
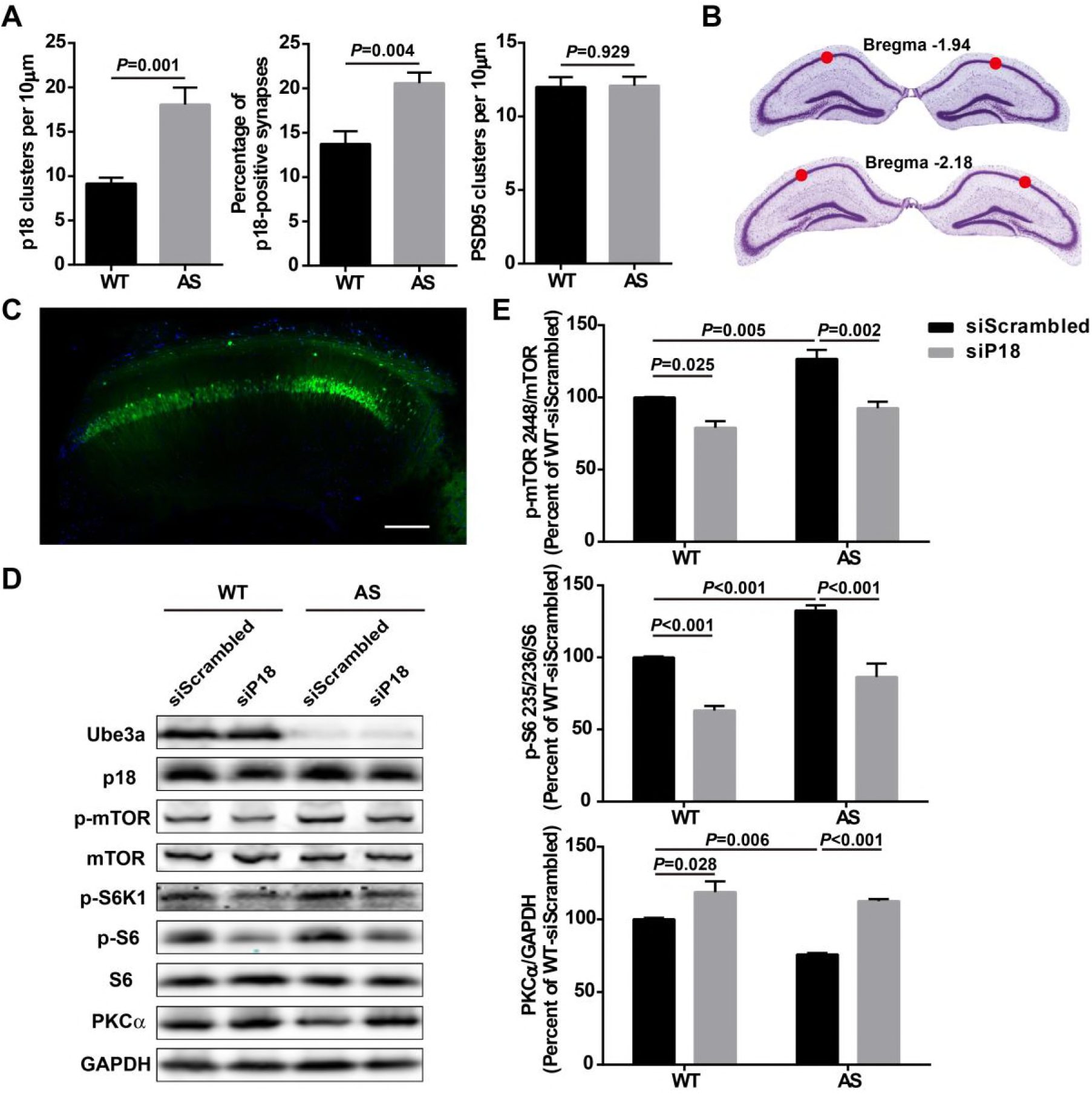
Effects of p18 knockdown in hippocampal CA1 region on mTOR signaling in WT and AS mice. Related to Figure 6. (A) Quantitative analysis of the number of p18- (left, p = 0.001) and PSD95-immunoreactive puncta (right, p = 0.929), as well as percentage of p18 and PSD95 dually stained puncta/synapses (middle, p = 0.004) in hippocampal CA1 region. N = 6, unpaired t-test. This data is paired with Figure 6A. (B) The coordinates of the injection sites were (mm): AP -1.94, ML ±1.4, DV -1.35 from Bregma; AP -2.2, ML ±1.8, DV -1.5 from Bregma, in the CA1 region of hippocampus and are indicated by red circles. (C) Representative tile scan confocal image of GFP expression in hippocampal CA1 region 4 weeks following injection of AAV with GFP reporter gene. Scale bar = 200 μm. (D) Representative images of Western blots labeled with Ube3a, p18, p-mTOR, mTOR, p-S6K1, p-S6, S6, and PKCα (GAPDH as a loading control). Protein lysates from hippocampal CA1 region infected with the indicated AAV were prepared for Western blot analysis. (E) Effects of p18 knockdown in hippocampal CA1 region on mTOR signaling in WT and AS mice. For p-mTOR, p = 0.025, WT-siScrambled vs WT-siP18, p = 0.005, WT-siScrambled vs AS-siScrambled, p = 0.002, AS-siScrambled vs AS-siP18; For p-S6, p < 0.001, WT-siScrambled vs WT-siP18, p < 0.001, WT-siScrambled vs AS-siScrambled, p < 0.001, AS-siScrambled vs AS-siP18; For PKC, p = 0.028, WT-siScrambled vs WT-siP18, p = 0.006, WT-siScrambled vs AS-siScrambled, p < 0.001, AS-siScrambled vs AS-siP18; n = 4 for WT-siScrambled, WT-siP18, and AS-siScrambled, n = 3 for AS-siP18, two-way ANOVA with Tukey’s post-test.

**Figure 6-figure supplement 2.**
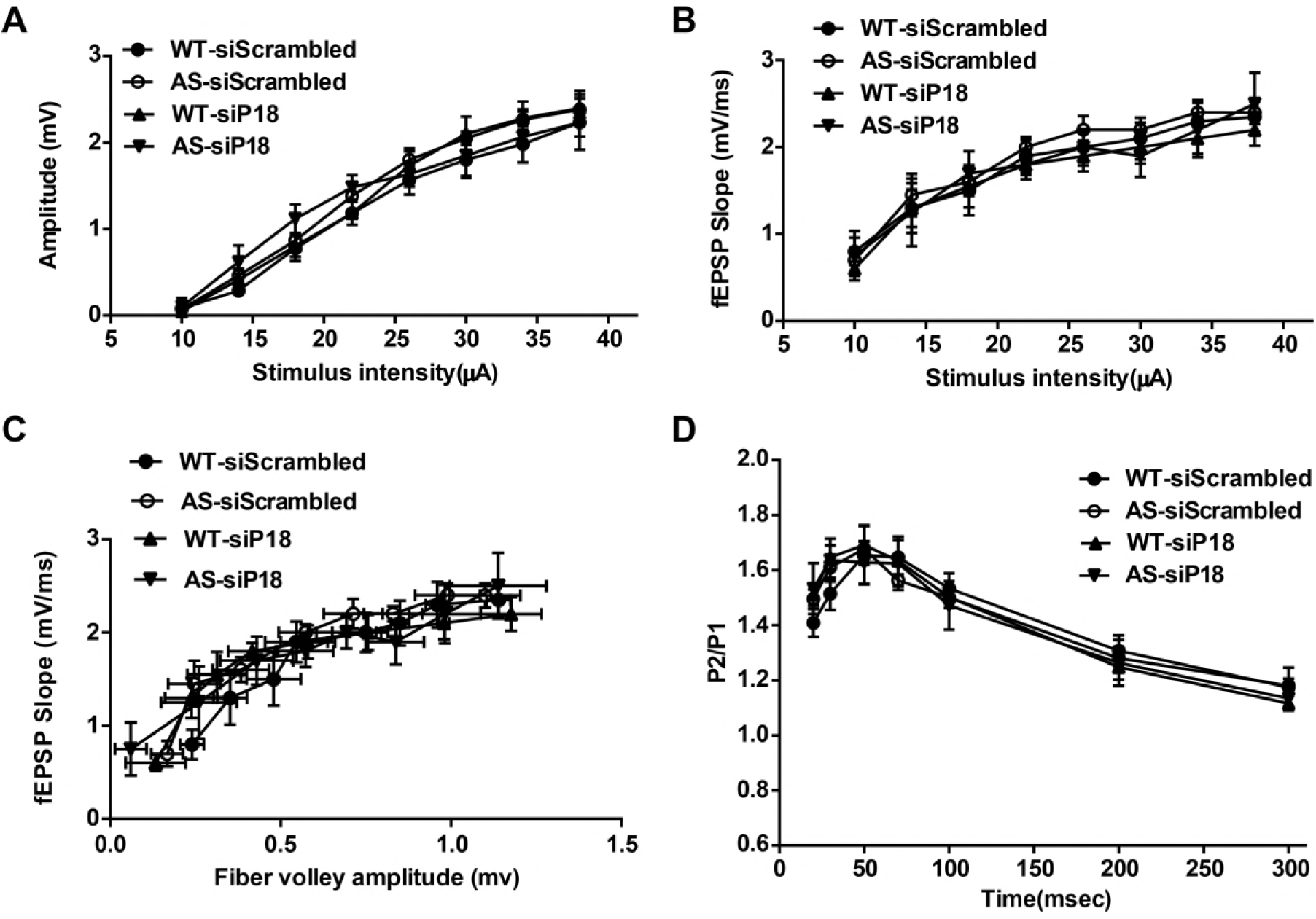
Effects of p18 knockdown in hippocampal CA1 region on input/output curves and paired-pulse facilitation in WT and AS mice. Related to Figure 6. (A-C) Input/output curves. Amplitudes of field EPSPs (A) and the slope of the field EPSP (B) were determined for various intensities of stimulation. (C) Relationship between the slope of the evoked fEPSPs and the corresponding fiber volley amplitude. The results are means ± S.E.M.; n = 3; there were no significant differences among the 4 groups of mice. (D) Paired-pulse facilitation. The amplitude of the second response of a paired-pulse was calculated as a percent of the amplitude of the first response for various inter-pulse intervals. The results are means ± S.E.M.; n = 5; there were no significant differences among the 4 groups of mice.

**Figure 7-figure supplement 1.**
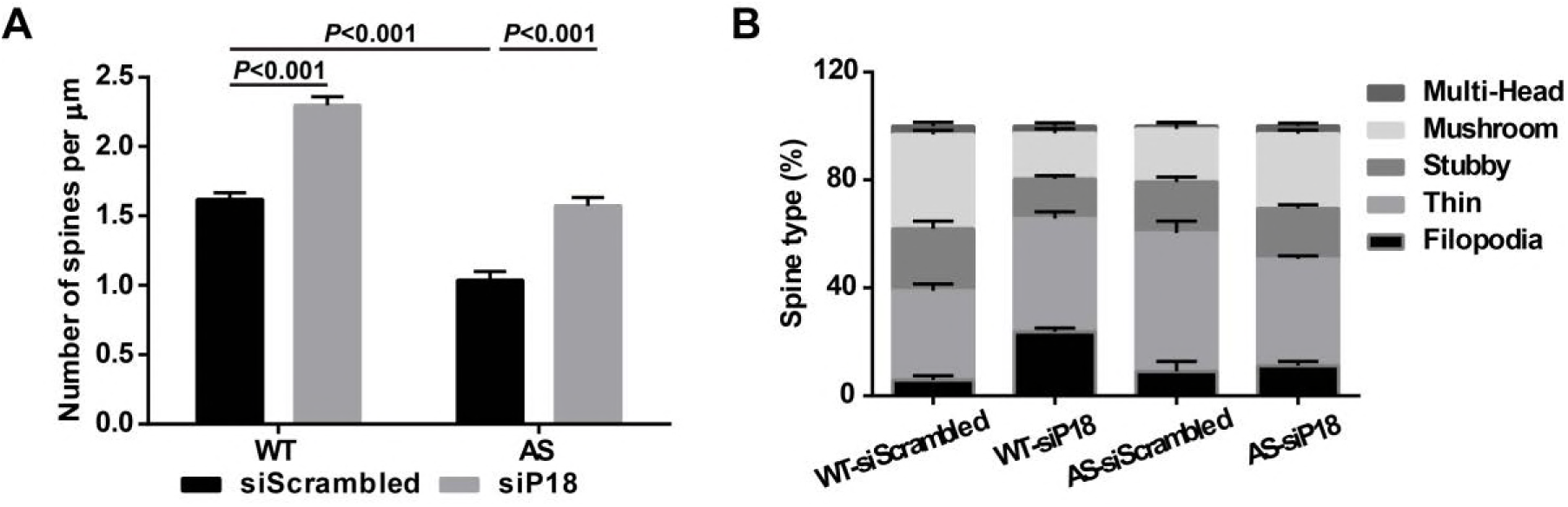
Effects of p18 knockdown in hippocampal CA1 region on dendritic spine number and the proportion of various spine types in WT and AS mice. Related to Figure 7. (A) Quantitative analysis of dendritic spine density shown in Figure 7A (means ± SEM from 10 slices). P < 0.001, two-way ANOVA with Tukey’s post-test. (B) Effects of p18 downregulation in hippocampal CA1 region on the proportion of various spine types in WT and AS mice.

